# Phage susceptibility to a minimal, modular synthetic CRISPR-Cas system in *Pseudomonas aeruginosa* is nutrient dependent

**DOI:** 10.1101/2025.01.14.632895

**Authors:** Josie F K Elliott, Bridget N J Watson, Keira Cozens, Edze R Westra, Tiffany B Taylor

## Abstract

CRISPR-Cas systems can provide adaptive, heritable immunity to their prokaryotic hosts against invading genetic material such as phages. It is clear that the importance of acquiring CRISPR-Cas immunity to anti-phage defence varies across environments, but it is less clear if and how this varies across different phages. To explore this, we created a synthetic, modular version of the type I-F CRISPR-Cas system of *Pseudomonas aeruginosa*. We used this synthetic system to test CRISPR-Cas interference against a panel of 13 diverse phages using engineered phage-targeting spacers. We observed complete protection against eight of these phages, both lytic and lysogenic and with a range of infectivity profiles. However, for two phages CRISPR-Cas interference was only partially protective in high nutrient conditions, yet completely protective in low nutrient conditions. This work demonstrates that nutrient conditions modulate the strength of CRISPR-Cas immunity and highlights the importance of environmental conditions when screening defence systems for their efficacy against various phages.

## Introduction

Our knowledge of the range of systems that prokaryotes use to defend themselves from the viruses that infect them – bacteriophage (phage) – has rapidly expanded in the last decade (Georjon & Bernheim, 2023). In conjunction with this, more high throughput studies are revealing the intricacies around which defence systems provide resistance to a host against which phage species (Gaborieau et al., 2023). Phages are highly diverse, such as recognising hosts through different receptors, carrying their own anti-defence systems, or replicating via different life cycles. Virulent phages undergoing a lytic lifecycle and temperate phages capable of either lytic or lysogenic lifecycles. In the lytic life cycle using host machinery to lyse and kill the cell, and in the lysogenic life cycle phage incorporating into the host genome (where they are called a prophage) to be replicated with the host genome, before excising to replicate and lyse the cell at some later time (P.-J. Ceyssens & Lavigne, 2010; Dion et al., 2020). Due to this diversity, no single defence system can offer protection to a host cell against all phages, and in turn it’s likely that no phage can evade every defence system. As interest in using phages to control bacterial growth in medical or agricultural settings grows (Brockhurst et al., 2021; Strathdee et al., 2023; Xu, 2021), an improved understanding of the traits enabling phages to evade bacterial defence systems, and whether any environmental factors can promote that evasion, will allow us to develop and apply more effective phage-based treatments.

One of the most well-studied bacterial defence systems is the CRISPR-Cas (clustered regularly interspaced short palindromic repeats – CRISPR associated) system (reviewed (Hille et al., 2018)). Upon entry to the cell, Cas (CRISPR-associated) proteins select, process and integrate regions of the invading DNA (called protospacers), into the CRISPR array (reviewed (Jackson et al., 2017)). Upon incorporation, the unique DNA sequences are called spacers, and the immunity they provide can be inherited. CRISPR arrays can contain many spacers, targeting the same or different mobile genetic elements (MGEs), and hosts can contain multiple CRISPR arrays (Weissman et al., 2018). This process of gaining new spacers is called adaption or spacer acquisition. Each spacer is transcribed and processed to produce a crRNA (CRISPR RNAs), which then form ribonucleoprotein complexes with Cas proteins. Through complementary base pairing, the spacer sequences guide Cas nucleases to degrade targeted genetic material, providing immunity to the host cell (Hille et al., 2018). This process of targeting, binding, and nuclease-mediated destruction of invading genetic material is termed interference.

In type I CRISPR-Cas systems acquisition of new spacer sequences into the CRISPR array can occur via either naïve or primed acquisition (Jackson et al., 2017). Naïve acquisition, is as described above, whereas primed acquisition occurs when there are pre-existing (priming) spacers, that may have complete or partial complementarity to the re-infecting foreign genetic material, these ‘priming’ spacers are able to enhance the acquisition of new spacers (Datsenko et al., 2012; Richter et al., 2014; Swarts et al., 2012).

Although in theory a CRISPR-Cas system could acquire spacers and target any incoming genetic element, the capabilities of CRISPR-Cas systems as phage defences are more curtailed in nature. Computational predictions have suggested that hosts with CRISPR-Cas systems may rarely outcompete those lacking the system during virulent phage epidemics, and during prophage epidemics hosts with CRISPR-Cas systems are only successful when the prophage incurs considerable cost to the host (Westra & Levin, 2020). Previous studies have navigated the challenge of natural acquisition of de novo CRISPR-Cas immunity by engineering species to insert spacers against phage to study functional aspects of the CRISPR-Cas system (Cui & Bikard, 2016; Jiang et al., 2013; Vorontsova et al., 2015). Interestingly, in these engineered strains it is observed that CRISPR-Cas systems do not always offer protection and benefit to the host. In *Escherichia coli* when the type I-E CRISPR-Cas system was directed against a variety of lytic phages, only spacers targeting pre-early genes (which transcribe and shut down host cellular functions before the entire phage genome enters) were able to elicit protection against the T5 phage. Against the T4 and R1-37 phages CRISPR-Cas was ineffectual at providing protection, due to an unknown CRISPR-Cas interference phage evasion mechanism (Strotskaya et al., 2017).

In some cases the inability of CRISPR-Cas systems to provide protection against phage lysis to their hosts can be explained by factors such as anti-CRISPR proteins (Acrs) that block CRISPR-Cas activity (Bondy-Denomy et al., 2013), or the proteinaceous nucleus-like structures some jumbo phage use to shield their genetic material from CRISPR-Cas systems (Malone et al., 2020), or alternatively the genetic material of phage can be disguised using epigenetic DNA modifications (Vlot et al., 2018). In other cases, CRISPR-Cas immunity has been suggested to be too slow to enable recovery from infection from virulent phage, resulting in an abortive infection phenotype (Watson et al., 2019). However, it is still unclear if the ecological advantage of a CRISPR-Cas system against any phage species may be predicted by traits of the phage.

Furthermore, it is yet unknown how factors that result in imperfect protection by CRISPR-Cas systems against phage replication may depend on ecology. The effect of environmental parameters on CRISPR-Cas evolution has been explored in previous work on *P. aeruginosa* UCBPP PA14 with its lytic phage DMS3vir phage (which is a mutant of the lysogenic phage DMS3 locked into the lytic lifestyle (Cady et al., 2012; Zegans et al., 2009)). The preference of populations to evolve CRISPR-Cas immunity via spacer acquisition, rather than via selection that enriches surface receptor mutants that prevent phage-binding (the type IV pilus in the case of DMS3vir), is highly sensitive to both biotic and abiotic factors. For example, the evolution of DMS3vir phage resistance via CRISPR-Cas spacer acquisition was favoured in low nutrient conditions (Westra et al., 2015), which may be due to increased mutational supply in high-growth conditions as well as selection for CRISPR immunity over surface mutants under low phage densities (Watson et al., 2023). Similarly, increased CRISPR-Cas adaptation was observed when PA14 populations were grown at lower than optimal temperatures (Høyland-Kroghsbo et al., 2018). Slowed growth in the presence of bacteriostatic antibiotics has also been shown to increase spacer acquisition and favour CRISPR-Cas immunity (Dimitriu et al., 2022). Similarly, translation inhibitory antibiotics increase CRISPR immunity levels against phage carry Acrs (Pons et al., 2023). This suggests that CRISPR-Cas systems may present a more formidable barrier to phage infection in limited resource environments. However, the *P. aeruginosa* model has only been monitored against DMS3vir or closely related phage (Broniewski et al., 2020), thus it is still unknown if the ecological sensitivity applies to a broader panel of phage. Additionally, it is unclear the influence ecological conditions may have on CRISPR-Cas interference only, as opposed to the evolution of immunity via spacer acquisition.

To address this we created a synthetic, minimal, version of the type I-F CRISPR-Cas system from *P. aeruginosa* made modular with restriction enzyme recognition sites for easy exchange of genetic elements such as spacers. The synthetic CRISPR-Cas system allowed us to engineer strains with complimentary spacers to a panel of phage with various characteristics and experimentally test whether CRISPR-Cas interference is equally effective against a diverse panel of phage, and whether the efficacy of CRISPR-Cas interference increases under low nutrient conditions.

## Materials and Methods

### Bacterial strains and phages

All strains cloned in this study were created from the genetically engineered ΔCRISPR-Cas version of the UCBPP-PA14 strain, described herein as ‘CRISPR-Cas knock-out’ (Cady & O’Toole, 2011). The synthetic CRISPR-Cas system was validated primarily using the phage DMS3vir, which was engineered from the phage DMS3 to be obligately lytic (Cady et al., 2012). During the efficiency of plating experiment the related phages were used: DMS3mvir-AcrIF1 (which contains a type I-F targeting anti-CRISPR gene) and DMS3mvir-AcrIE3 (Bondy-Denomy et al., 2013). The CRISPR-Cas system was screened against the *P. aeruginosa* temperate/lysogenic phages detailed in table S1.

All overnight cultures were incubated at 37°C shaking at 200 r.p.m in LB (Miller’s lysogeny broth) or M9 medium (22 mM Na_2_HPO_4_; 22 mM KH_2_PO_4_; 8.6 mM NaCl; 20 mM NH_4_Cl; 1 m MMgSO_4_; 0.1 mM CaCl_2_) supplemented with 0.2% glucose.

### Cloning synthetic CRISPR-Cas system into *P. aeruginosa* using mini-Tn7 insertion

The synthetic CRISPR-Cas DNA, ordered from GeneArt Thermofisher Scientific (https://benchling.com/s/seq-gNNEFhMwdBo2URdFk7Y8), was based on the *P. aeruginosa* PA14 type I-F CRISPR-Cas system, minimized for functionality with restriction sites added for modularity (fig 1) The CRISPR array contains a single repeat-spacer-repeat sequence from the CRISPR 2 array (shown to acquire new spacers more frequently (Heussler et al., 2016; Høyland-Kroghsbo et al., 2018)). The leader sequence of the CRISPR 2 array was annotated as the 134 base pairs upstream of the first CRISPR repeat (sequence supplied by Dr Vorontsova (Vorontsova et al., 2015)). The promoter was assumed to be contained within 135 bases upstream of the leader sequence and downstream of the adjacent gene *PA14-3370*. The non-targeting spacer, with two BbsI recognition sites and a MluI site for modularity, was designed that had no targets in the *E. coli* or *P. aeruginosa* chromosomes. A strong rrnB T1 terminator was added downstream to prevent read-through from the CRISPR array (Noirot-Gros et al., 2018), flanked by NsiI restriction sites. The *cas* operon promoter, assumed to be between the CRISPR 2 array and *cas1* genes, was made interchangeable with NotI and NcoI sites. Other restriction enzyme recognition sites (BbsI, MluI, NsiI, NotI, NcoI and AscI) were then edited out using *P. aeruginosa* codon usage tables (Nakamura et al., 2000; West & Iglewski, 1988).

**Figure 1.**
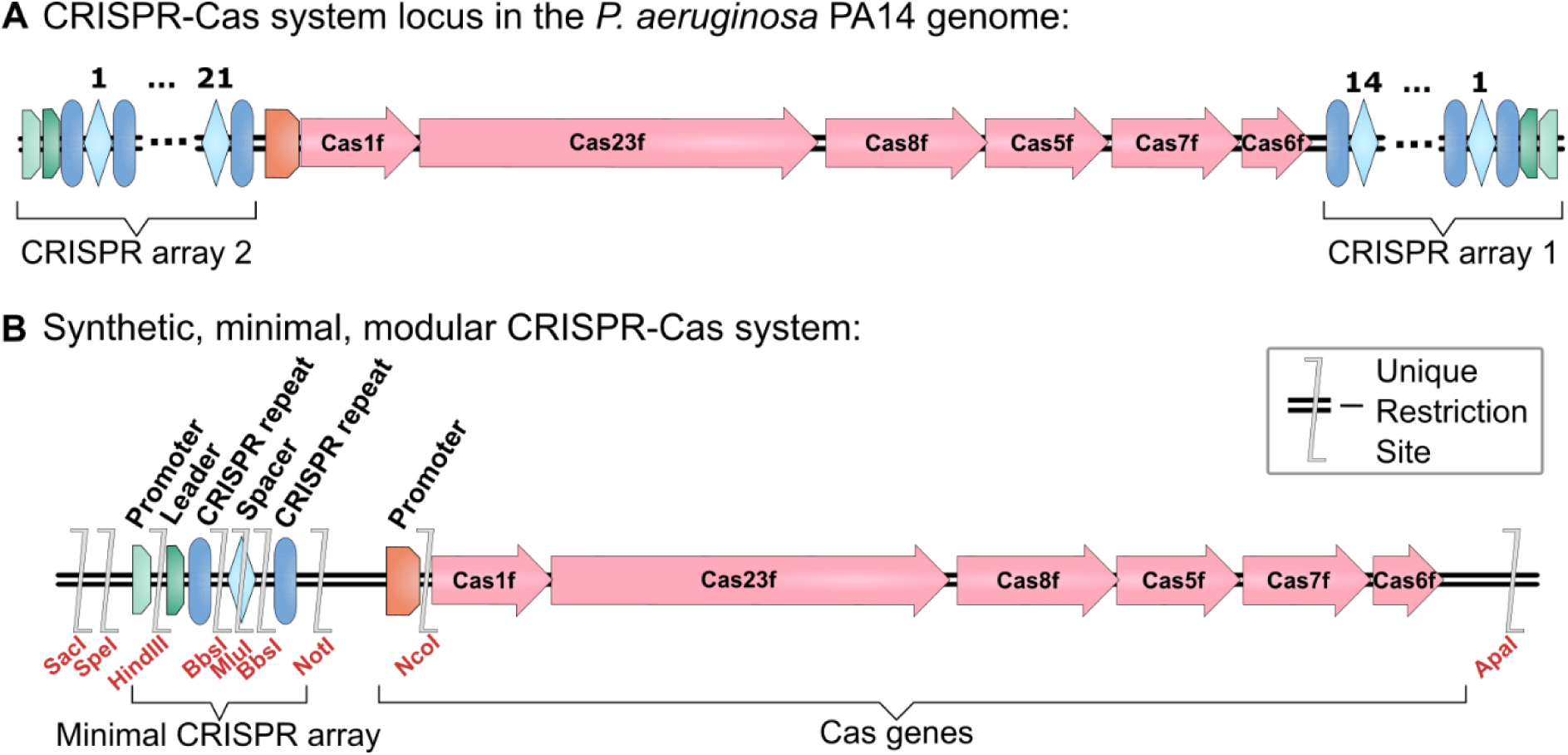
– **Validation of a synthetic version of the type I-F CRISPR-Cas system of *P. aeruginosa* PA14**. (A) Schematic of the CRISPR-Cas system locus in the genome of *P. aeruginosa* PA14 (adapted from Cady et al., 2012). This contains two CRISPR arrays, and 6 *cas* genes: Cas1f – endonuclease Cas1, Cas23f – helicase also known as Cas3, Cas8f – also known as Csy1, Cas5f – also known as Csy2, Cas7f – also known as Csy3, Cas6f – endoribonuclease also known as Cas6/Csy4 (names updated from the Cady et al., 2012 in line with naming conventions in Makarova et al., 2019). In the genome this locus in antisense but has been displayed in the same sense orientation as the synthetic CRISPR-Cas system for ease of comparison. The relative lengths of each gene in the schematic have been drawn proportional to sequence length. (B) Schematic of the synthetic, minimal, modular CRISPR-Cas system which has the CRISPR arrays reduced to one repeat-spacer-repeat from the CRISPR 2 array. The relative positions of the added restriction sites are shown.

The synthetic CRISPR-Cas DNA was transferred into the pUC18T-mini-Tn7T-Gm plasmid (pUC18T-mini-Tn7T-Gm was a gift from Herbert Schweizer - Addgene plasmid # 63121 ; http://n2t.net/addgene:63121 ; RRID:Addgene_63121)(Choi & Schweizer, 2006) via restriction-ligation cloning using ApaI and SpeI (New England Biosciences). To incorporate new spacers against either plasmid or phage, the spacer sequences were ordered as two overlapping primers which, when annealed, create sticky ends compatible with the BbsI restriction sites. Oligonucleotides were annealed by heating in annealing buffer (100 mM potassium acetate, 30 mM HEPES, pH 7.5) at 95 °C for 5 minutes before slowly cooling (30 seconds 0.5 °C stepwise cooling to room temperature). The spacer DNA was phosphorylated with T4 Polynucleotide Kinase before ligation with the pUC18T-mini-Tn7T-Gm-CRISPR-Cas plasmid (digested with BbsI and dephosphorylated with Shrimp Alkaline Phosphatase (New England Biosciences)). Successful insertion was determined by PCR with a forward primer against the pUC18T-mini-Tn7T-GM plasmid (Tn7-F, table S2) and a reverse primer after the CRISPR array (CRISPR-R, table S2), then confirmed via sequencing (Sanger sequencing, Source Bioscience).

The synthetic CRISPR-Cas DNA was inserted as a single copy into the chromosome of *P. aeruginosa* CRISPR-Cas knock-out using the four-parent conjugal puddle mating conjugation version of the protocol from Choi & Schweizer, 2006. PCR (glmS-F and CRISPR-R primers, table S2) was used to verify insertion.

To create phage targeting spacer strains, oligonucleotides of two spacers (separated by a repeat sequence), targeting different regions in the phage genome were cloned as described above into the synthetic CRISPR-Cas system (table S3). Spacer 1 targeted gene regions near phage genome ends (and these regions were verified by PCR and sanger sequencing of the phage), while spacer 2 targeted central gene sequences. This was based on research identifying the linear genome ends as protospacer sampling hotspots (Levy et al., 2015).

### One step phage growth curves

*P. aeruginosa* wildtype cultures were grown overnight in LB then adjusted to OD_600_ of 2.6 in 1 ml LB and inoculated into 25 ml culture of LB in a 250 ml flask to give a starting OD_600_ of 0.1. Cells were grown shaking at 37 °C to early exponential phase (OD_600_ 0.2-0.3) at which point 100 μl of phage at ∼10^9^ (pfu/ml) was added. This gave a starting phage titre of ∼4 x 10^6^ pfu/ml and starting multiplicity of infection (MOI) of ∼ 0.01. Samples were taken at various time points up to 120 min post infection and added to PBS and chloroform (sample:chloroform 10:1 v/v) to lyse cells, allowing measurement of total number of mature phages. Samples were serially diluted and 2 μl was spotted in triplicate on a lawn of wildtype *P. aeruginosa* PA14. The phage PA14P2 had to be grown and plated on *P. aeruginosa* PA14 *csy3::LacZ* as the wildtype CRISPR array carries a spacer against this phage. Phage burst size was calculated from the number of phages released as the maxima of the growth curve (max pfu/ml – min pfu/ml) / the number of cells infected (t=0 pfu/ml – min pfu/ml). Phage adsorption over time was calculated as the phages lost from the media to the minima of the growth curve (t=0 pfu/ml – min pfu/ml)/(t=0 pfu/ml). The latent period was defined at the minima of the growth curve before the phage burst starts.

### Phage virulence assays

The method to determine phage virulence was adapted from the metrics in Storms et al., 2020. Overnight cultures of wild type *P. aeruginosa* (and *csy3::lacZ* for PA14P2 phage) were grown in 10 ml LB then normalised to OD_600_ of 1. Each culture was resuspended in PBS and added to sterile LB in 96-well plates at 1:100 dilution and growth curves were performed in biological and technical triplicate. Phages were diluted from the highest harvestable titre (to a max of 10^9^ pfu/ml) by a factor of 10 across 5 dilutions and added 1:100 into the wells as well as a PBS no-phage control. Phage and bacterial titre were also plated and measured at the start of the experiment to allow calculation of accurate multiplicity of infection (MOI). OD_600_ measurements were taken every 10 minutes for 23 hours while the plate was incubated at 37 °C shaking at 180 r.p.m.

To prevent the selection for phage resistant cells impacting the calculated virulence index, the growth curve measurements were cut off at the time at which the average OD readings for the samples with the highest concentration of phage surpassed the average OD readings for the second highest concentration of phage (or the third highest if the first two concentration could not be resolved). Local virulence was calculated as 1 - (area under the curve at each given MOI / area under the curve at no phage dilution). Virulence index was then calculated from local virulence as the area under the local virulence curve divided by the theoretical maximum area under the virulence curve (area if local virulence equalled 1 for all MOI).

### Bacterial growth curves

Overnight cultures of each strain were grown in 10 ml of LB then normalised to OD_600_ of 1. Each culture was resuspended in PBS and added a 1:100 dilution to LB or M9 in sterile 96-well plates. Phage cultures were diluted in PBS and added at 1:100 to the wells to give each concentration. OD_595_ measurements were then taken every 10 minutes for 23 hours while the plate was incubated at 37 °C shaking at 180 r.p.m. Phages were added at two high (10^8^ pfu/ml diluted 1:100 into 96-well plate), low (10^6^ pfu/ml diluted 1:100 into 96-well plate) concentrations, as well as PBS being added for the no-phage control.

### Statistics

All statistical analysis were performed in the R statistical package 4.4.0 (2024-04-24)(R Core Team, 2024). Data that ranged over three powers of ten or more were log transformed to normalise residuals. Statistical significance for the efficiency of plating and conjugation efficiency, was determined using one-way ANOVA analysis with post-hoc analysis via Tukey test. Pairwise comparisons depicted on graphs were t-tests, any p-value adjustments being detailed in figure legends.

## Results

### A minimal, modular, synthetic CRISPR-Cas system allows for a functional, genetically tractable defence system in the host genome

We sought to create a version of a CRISPR-Cas system within which all the components can be easily edited. This would be a potentially useful tool for studying how the mechanisms of CRISPR-Cas systems impact co-evolution with phages. The type I-F CRISPR-Cas system of *P. aeruginosa* UCBPP-PA14 contains two CRISPR arrays (CRISPR 1 and CRISPR 2) which flank six *cas* genes (Cady et al., 2012; Makarova et al., 2020) (fig 1A). This spans around 14kb in the genome. A synthetic version of this CRISPR-Cas system was designed to be minimal, modular, and genetically tractable (fig 1B). The two CRISPR arrays were reduced to a single array with a single spacer flanked by repeats. The system is also made modular due to the addition of restriction endonuclease (RE) recognition sites flanking key regions (fig 2B). The type-II RE recognition sites within the spacer face outwards, allowing scarless replacement of the spacer with any number or design of spacers using either short oligonucleotides with simple restriction ligation cloning, or multiple fragments using golden gate cloning as has been used to produce multi-spacer guide RNAs (Shaw et al., 2022). Thus, the synthetic CRISPR-Cas system can be quickly and easily edited to target any required sequence that has suitable protospacer adjacent motif recognition sites.

This synthetic CRISPR-Cas system was stably integrated as a single copy into the host chromosome using a Tn7 transposon-based method, eliminating the need for plasmid maintenance and consistent antibiotic selection, thus making it ideal for long-term evolution experiments. Furthermore, this avoids dosage variability that may be associated with plasmid copy number. The insertion of the synthetic CRISPR-Cas system into the *P. aeruginosa* CRISPR-Cas knock-out strain did not result in a fitness cost to the host (fig S1A) and includes a selectable gentamicin resistance marker for easy identification of edited strains. The synthetic CRISPR-Cas system was shown to be capable of specifically targeting plasmids (fig S1B), was able to target the DMS3vir phage at similar efficiencies as the native CRISPR-Cas system (fig S1C), and was shown to be capable of acquiring spacers via both primed and naïve acquisition (fig S1D). Taken together these results show that the synthetic CRISPR-Cas system retains all activities of the native CRISPR-Cas system when complemented into *P. aeruginosa*.

### The panel of phage used to test CRISPR-Cas interference efficacy possessed a broad range of infectivity metrics

We initially hypothesised that the efficacy of CRISPR-Cas interference may vary with traits associated with phage biology such a virulence, e.g. CRISPR-Cas immunity may be more effective against less virulent phage. We therefore selected a panel of 13 diverse dsDNA phages (dsDNA being the most sampled phage type) including those with lytic and lysogenic lifecycles, and that use different surface receptors, to test whether CRISPR-Cas interference and ability to provide protection against phages is dependent on any predictable characteristics of a phage (fig S2). These phages were DMS3vir, DMS3, JBD5, JBD10, JBD88b, PA14P1, PA14P2, LMA2, JBD63c, LBP1, JBD18, JBD25, phiKZ. To quantify the characteristics of these phages we performed one step growth curve, and phage virulence experiments (fig 2).

**Figure 2.**
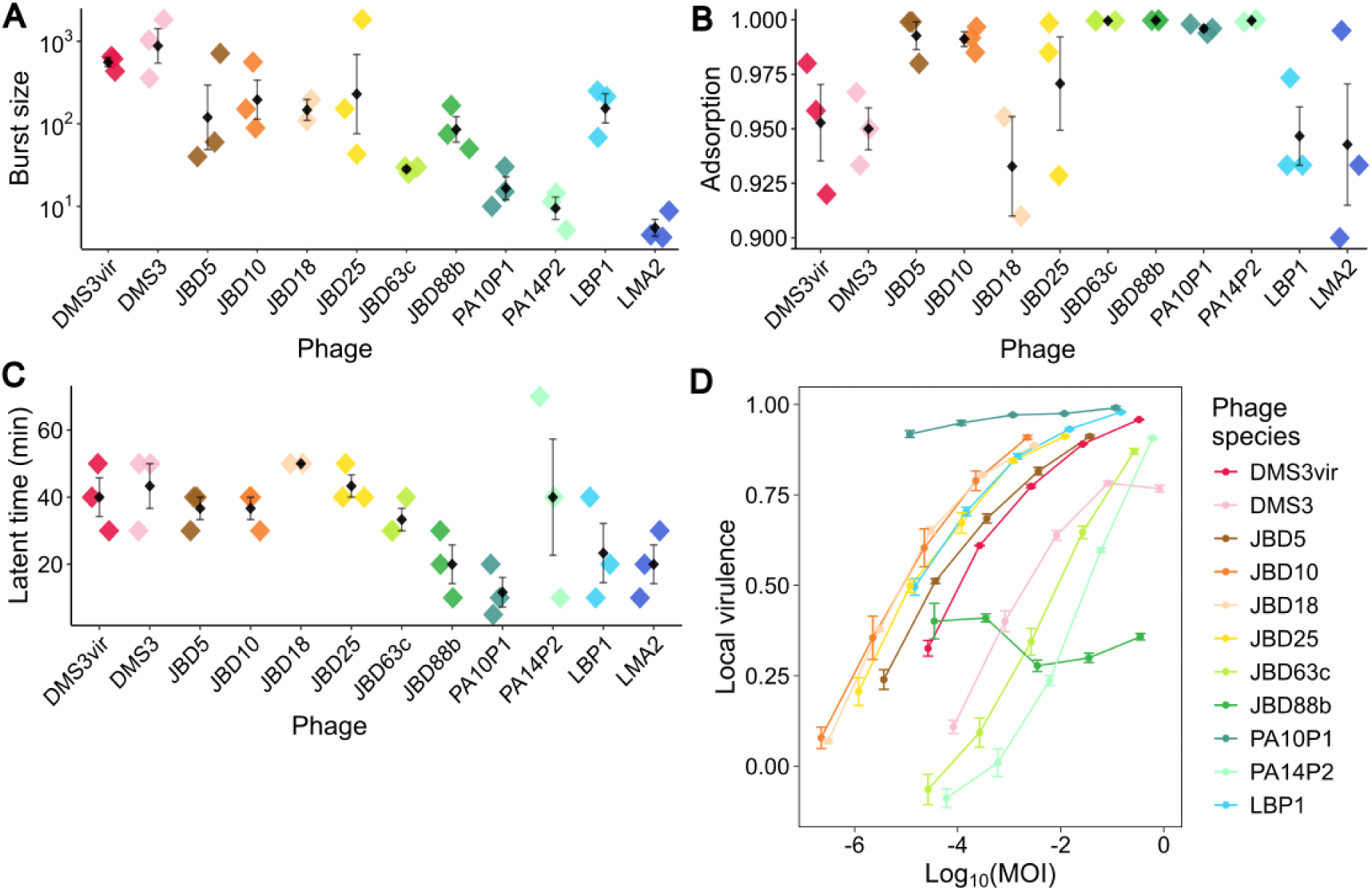
– **Comparison of the infection characteristics of a panel of 13 phages**. Each phage is presented in the same colour across each panel. Assays were performed on wildtype *P. aeruginosa* strains (apart from PA14P2 phage which used a *csy3::lacZ* knock-out strain). (A-C) Black diamonds represent mean values (N=3) and error bars show the standard error around the mean. Diamonds coloured by phage show individual replicates (A) Burst size represents the number of phages released per cell during the growth curve during the burst phase of the one step growth curve when the first mature phages are assembled post infection. (B) Adsorption describes the fraction of total phage population taken up by the cells in the media during the first part of the growth cycle. (C) Latent time describes the time taken from initial infection to the when the first mature phages start to be assembled and subsequently released into the media. (D) Plot of local virulence across log multiplicity of infection (MOI). Circles represent mean values (N=3), and error bars show standard error around the mean. Local virulence represents the area under a growth curve at each MOI of phage infection, normalised against the growth curve when no phages are present.

The one step growth curve assay standardises the start of the phage infection cycle for all phages in a sample allowing measurements of how many phages are released per cell (burst size, fig 2A), what fraction of the phage population successfully adsorb to and inject their DNA into cells (adsorption, fig 2B), and how long the phage replication cycle until mature phages are assembled (latent time fig 2C). The plaques from jumbo phage phiKZ on solid media were not able to be resolved due to their hazy morphology, thus phiKZ was excluded from this experiment. The remaining phages showed a range of phenotypes that vary between infection metrics, e.g. the phage with the highest burst size – DMS3 – does not have the highest adsorption or shortest latent time. Each individual growth curve is shown in supplementary figures S4-S16A. Together this panel of phage thus able to represent a range of infectivity metrics, allowing us to test CRISPR-Cas interference against phage with a range of lifestyles, receptors, and virulence.

The ability of each phage to infect and prevent growth of the host bacteria was assessed using a virulence assay in liquid culture (fig 3D). In this assay, phages are diluted to give a range of phage-to-cell ratios values and the effect on growth measured by normalising the area under the growth curve at each phage dose to the area under the growth curve in the absence of phages, which produces the metric ‘local virulence’. Individual growth curves at each phage dose are shown in supplementary figures S4-S16B. The area under the local virulence curve can be also calculated and normalised against the maximal possible area to give an individual value termed virulence index (fig S2). The higher the local virulence the more virulent the phage, with a maximum value of 1 if there is no growth in the presence of the phage. PhiKZ was excluded as it could not be grown to a high enough titre to give at least three log_10_ dilutions that impacted growth. The LMA2 phage was also excluded as it elicits bacterial precipitate when cultured in liquid, which interfered with the spectrophotometer OD_595_ readings. Of the eleven phages tested, there is a clear diversity in phage virulence from the most virulent PA14P2 to the least virulent PA10P1 and JBD88b.

We were able to observe that the phenotypic diversity of the phage panel was also reflected in the genetic diversity of the phage used. A phylogenetic tree of all complete phage genomes labelled as targeting *P. aeruginosa* in the NCBI database (fig S3) demonstrated that phylogenetic relatedness between phages could not explain patterns of virulence. Trees were constructed based on the amino acid sequence of both the terminase large subunit protein (fig S3A) and the major capsid protein (fig S3B) to ensure all 13 phages in our panel were included.

### CRISPR-Cas can provide complete protection against most phages but nutrient-dependent protection against others

The ease with which the spacer in the synthetic CRISPR-Cas system can be interchanged with any phage targeting sequence desired presented an opportunity to test CRISPR-Cas interference against each of the 13 phages. Synthetic CRISPR-Cas *P. aeruginosa* strains were created that had two spacers designed to target different regions of the genome in each phage (table S3). This ensured that the ability of the phages to evolve escape mutations against both spacers over the experimental period and confound results would be near impossible. Each spacer 1 in the phage-targeting strain CRISPR-Cas array was also validated by testing the ability of the CRISPR-Cas system to reduce conjugation efficiency when a short region of the phage genome containing the protospacer was cloned into a plasmid (fig S4-11D). These phage-targeting synthetic CRISPR-Cas strains were then tested to see if they could restore normal growth in the presence of each phage. The synthetic CRISPR-Cas system was able to provide complete immunity to eight of the phages tested: restoring the growth in liquid high nutrient media to similar levels as when phages were absent (DMS3vir, DMS3, JBD5, JBD10, JBD88b, PA10P1, PA14P2 and LMA2 (fig S4-11E)).

Example data for the DMS3vir phage is given in figure 3A. For each phage, this phenotype was also confirmed on solid media using an efficiency of plating assay (fig S4-11C). This use of the synthetic CRISPR-Cas system demonstrates that interference by the CRISPR-Cas machinery can provide protection against both lytic and lysogenic phages, across phages with a range of virulence strength, that target different receptors, and against phages with a range of infection dynamics such as adsorption time, burst size and latent time (fig S3).

**Figure 3.**
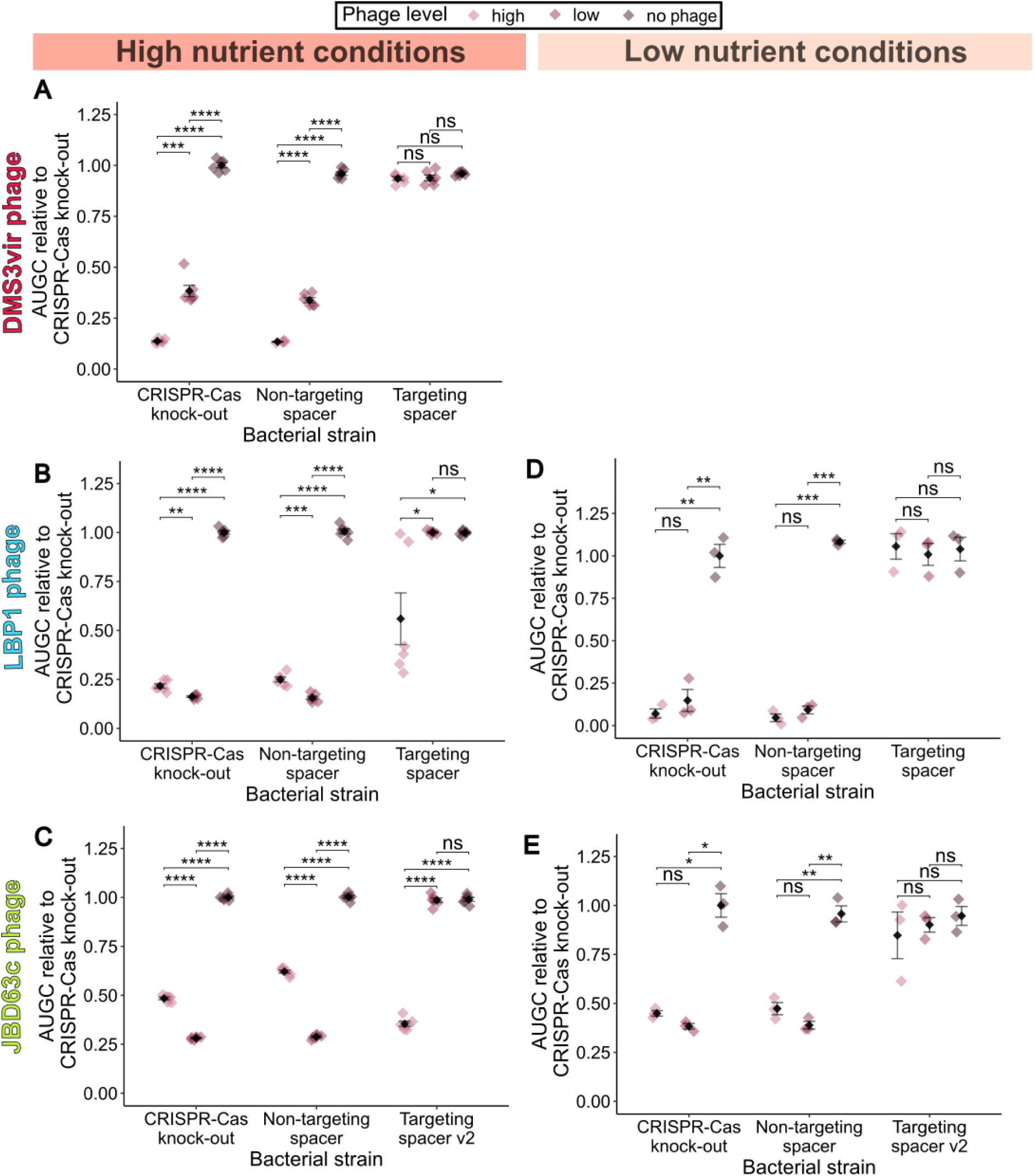
**– Growth dynamics in response to the various phage with and without a CRISPR-Cas system**. Area under growth curve (AUGC) was normalised relative to the average area under the growth curve for the CRISPR-Cas knock-out strain without phages added and plotted for each strain. The black diamond shows the mean (N=6 for high nutrient conditions experiments and N=3 for low nutrient condition experiments) for each strain with error bars representing the standard error around the mean. Diamonds for each individual datapoint are coloured by the phage level. Within each strain, the means at each phage level were compared with t-tests with FDR (false discovery rate) p-correction (* p<0.05, ** p< 0.01, *** p<0.001, **** p<0.0001, ns non-significant). (A) AUGC plot against bacterial strain with high, low or no DMS3vir phage added in high nutrient conditions. (B) AUGC plot against bacterial strain with high, low or no LBP1 phage added in high nutrient conditions. (C) AUGC plot against bacterial strain with high, low or no LBP1 phage added in low nutrient conditions. (D) AUGC plot against bacterial strain with high, low or no JBD63c phage added in high nutrient conditions. (E) AUGC plot against bacterial strain with high, low or no JBD63c phage added in low nutrient conditions.

For two phages, LBP1 and JBD63c, CRISPR-Cas interference with a phage targeting spacer only produced an ‘intermediate’ protective phenotype (fig 3B and fig 3C respectively). For the strains of *P. aeruginosa* with a CRISPR-Cas system containing a phage targeting spacer, in the presence of low levels of either LBP1 phage or JBD63c phage, the CRISPR-Cas system was able to restore bacterial growth levels to equivalent growth in the absence of phage. However, at high levels of phage, growth levels were reduced. In solid media experiments, the CRISPR-Cas system was able to produce a significant, but small in magnitude, decrease in efficiency of plating (fig S12-13C). The genomes of both phages possessed predicted hits for anti-CRISPR genes, although this was not unique to these two phages (fig S2).

Slowed growth, such as through reduced nutrient concentrations, has previously been shown to favour the evolution of CRISPR-Cas in *P. aeruginosa* (Westra et al., 2015), we repeated these liquid growth experiments in low nutrient media. In the presence of both LBP1 and JBD63c, strains with a phage-targeting spacer were then able to provide complete immunity in the presence of both low and high phage levels (LBP1 fig 3D, JBD63c fig 3E).

For the five phages where the initial phage targeting strains were not able to restore normal growth in nutrient rich media the presence of phages – LBP1, JBD63c, JBD18, JBD25 and phiKZ – a second version of the phage-targeting strain was created to ensure lack of complete phage defence wasn’t due to poor spacer choice. These second version spacers targeted new regions on the phage genome. CRISPR-Cas systems were tested via the same assays for ability to provide immunity to the host. For all these spacers that did not successfully target phage, the spacers in the synthetic CRISPR-Cas system were still able to successfully target the protospacer DNA when it was instead incorporated into a plasmid (fig S12-16D). This showed that the observed phenotype was due to some aspect of the phage, rather than anything intrinsic to the selected spacer sequence.

Interestingly, for the JBD63c phage, one phage targeting spacer strain did not provide any protection, however for the second version of the phage targeting strain that used a different spacer sequence, the CRISPR-Cas system was able to provide intermediate protection in high nutrient conditions and complete protection in low nutrient conditions (fig S13 E-F). This shows that spacer choice can also impact the success of CRISPR-Cas interference. The phages JBD18 and JBD25 showed complete resistance to CRISPR-Cas interference, including when experiments were repeated with new phage targeting spacers and in low nutrient conditions (fig S13E-F, fig S14E-F respectively), despite these spacers successfully targeting plasmids (fig S13-14D). However, given the demonstration by the JBD63c phage that successful CRISPR-Cas interference can be spacer-dependent, and that only two spacer variations of the synthetic CRISPR-Cas system in *P. aeruginosa* were tested, it remains likely that another spacer could yield immunity to these phages. The phage phiKZ also showed complete resistance to CRISPR-Cas interference irrespective of alternate spacers or changing nutrient conditions (fig S16). This was expected as phiKZ is a jumbo phage that builds a proteinaceous nucleus like structure around its DNA (Krylov et al., 2021), such structures have been shown to make the phage resistance to type I-F CRISPR-Cas systems (Malone et al., 2020).

Together these results show that upon acquiring spacers against a phage, CRISPR-Cas systems can provide robust immunity against a wide variety of phages. However, interference is not perfect, being modulated by the choice of acquired spacer (placing another barrier to successful immunity if not only a cell must acquire new spacers, but those spacers can only be from certain regions of the phage genome). Furthermore, we show for the first time that interference can be modulated by environmental conditions.

## Discussion

CRISPR-Cas systems are unique in their ability to provide adaptive, heritable immunity to their hosts by the acquisition of spacers derived from invading genetic material paired with the sequence-guided destruction of matching DNA through nuclease-mediated interference. Although clearly offering an advantage, CRISPR is not ubiquitously distributed across bacterial genomes (Tesson et al., 2022). Furthermore, the evolution of phage protection via mutation of surface receptors targeted by phages is frequently shown to be favoured in populations over the evolution of CRISPR-Cas immunity via spacer acquisition (Gurney et al., 2019; Westra et al., 2015). As the repertoire of known defence systems, and subtypes of these defence systems expands, there is increasing interest in an ability to map and predict which defences are robust against which phage (Gaborieau et al., 2023; Georjon & Bernheim, 2023) and this will be important in the development of ‘evolution-proof’ phage-based therapeutics (Brockhurst et al., 2021).

Understanding the molecular details of phage and CRISPR-Cas system co-evolution will be aided by genetic tools that allow for the precise dissection of the molecular components of the CRISPR-Cas system. To address this, we adapted the type I-F CRISPR-Cas system of *P. aeruginosa* to create a synthetic, minimal, modular CRISPR-Cas system where each element can be replaced by restriction-ligation cloning in *E. coli* and the whole system can then be stably integrated into the host genome using a highly repeatable Tn7 transposon-based system (Choi & Schweizer, 2006). For example, strains with any desired spacer sequence or number of spacers can be created with inexpensive oligonucleotides in 1-2 weeks (Shaw et al., 2022), as well as the potential to also change promoter sequences or integrate other genetic sequences alongside the CRISPR-Cas system. This contrasts with previous studies where CRISPR-Cas systems have had to be exposed to plasmids (Fineran et al., 2014), phages (Guillemet et al., 2022), or libraries of phage fragments (Strotskaya et al., 2017) in plasmids, from which acquisition events are detected and selected for. Other studies which have probed the molecular mechanisms of the *P. aeruginosa* type I-F CRISPR-Cas system have hosted *cas* genes and CRISPR arrays on plasmids (Vorontsova et al., 2015). The mini-Tn7 transposon-based integration system allows the CRISPR-Cas system to be stably inserted in the genome at a site-specific intergenic region, thus removing the requirement for continual selection for plasmid maintenance, and the potential effect of plasmid dosage as an extraneous variable. The Tn7 system can be applied to any host with an *att-Tn7* site in the genome (Choi & Schweizer, 2006; Gerth et al., 2012) making this system applicable for integration of the synthetic CRISPR-Cas system to multiple potential species.

After integrating the synthetic CRISPR-Cas system into a strain of *P. aeruginosa* PA14 that had its native CRISPR-Cas deleted, and then confirming the synthetic system was fully functional against plasmids and the DMS3vir phage (fig S1), we then created strains with two spacers each that both targeted one of 13 different phages. The panel of phages represented phylogenetically diverse origins, had a range of traits (being lytic and lysogenic), targeted different receptors, and displayed a range of values of infection dynamics (burst size, adsorption, latent time) and virulence (fig 2). This mirrors previous studies that have tested *P. aeruginosa* strains on a range of phages to investigate the evolution of spontaneous resistance mutations (Markwitz et al., 2022; Wright et al., 2018). However, as we engineered each CRISPR-Cas containing *P. aeruginosa* strain to already possess spacers against the phage, we were able to study the efficacy of CRISPR-Cas interference in isolation, unbiased by the likelihood of prerequisite spacer acquisition.

For eight of the 13 phages tested, CRISPR-Cas interference was able to provide complete protection of growth in liquid culture and significant decreases in titre when plating phages on lawns of each strain (fig 3, fig S4-11). This demonstrates that the measured phage characteristics could not be used to predict whether a CRISPR-Cas system could provide robust phage interference and protection to the host. For the lysogenic LBP1 (fig 3, fig S13) and JBD63c (fig 3, fig S12) phages, CRISPR-Cas interference provided intermediate protection in high nutrient liquid growth conditions, but complete protection in low nutrient conditions, thus showing that environmental parameters can affect the functioning of CRISPR-Cas interference.

Several previous studies have observed interactions between phage infectivity and nutrient conditions. For example, a study in *E. coli* investigating the maintenance of lytic lambda phage population in the presence of resistant cultures found that phages were only maintained in seven of twelve replicate populations in maltose-limited minimal media but maintained in all replicates grown in high nutrient both (Chaudhry et al., 2018). However in a single cell microfluidics experiment with *E.coli* and its phage T4, it was observed that increasing nutrients increased collective bacterial survival in the presence of phages (Attrill et al., 2023).

Other work has suggested a link between growth rate and the evolution of CRISPR-Cas immunity via spacer acquisition. When populations of *P. aeruginosa* PA14 were cultured with DMS3vir in the presence of bacteriostatic antibiotics, spacer acquisition increased and evolution CRISPR-Cas immunity was favoured over surface-modification based resistance (Dimitriu et al., 2022). This preference towards CRISPR-Cas immunity was also observed with growing the bacteria on carbon sources upon which growth was slowed (Dimitriu et al., 2022). This is in line with previous work showing that when grown in nutrient limited media, evolution of phage resistance via CRISPR-Cas immunity is favoured over surface-modification based resistance (Westra et al., 2015). Slower growth through reduced temperature was shown to promote both CRISPR-Cas adaption and interference in *P. aeruginosa* PA14, including upregulation of the Cas complex (Høyland-Kroghsbo et al., 2018). These results seem to be general across bacteria; in *Pectobacterium atrosepticum cas* gene expression was shown to increase upon deletion of the *galK* gene involved in galactose metabolism, suggesting a role for metabolic status in CRISPR-Cas regulation (Hampton et al., 2019).

Furthermore, phage dose and environmental conditions can impact the efficacy of Acr genes possessed by a phage. Broadly the pattern of predicted Acr genes in the genomes of each phage does not correlate with the observed patterns of successful CRISPR-Cas interference (fig S2), although these genes are predicted in both LBP1 and JBD63c. However, different Acrs only interfere with specific CRISPR-Cas system types. Previous studies have shown that phage in possession of Acr genes are able to proliferate in CRISPR-Cas carrying hosts at high phage titres but not at low phage titres due to phage cooperation (Landsberger et al., 2018). In addition to this, the addition of bacteriostatic antibiotics the environment slows the translation of phage genes, enhancing the protection CRISPR-Cas systems are able to provide to the host against phage with Acrs (Pons et al., 2023).

Although both versions of the phage targeting spacers against the JBD63c phage were functional when the protospacer was incorporated into a plasmid (fig S12D), one version of the phage targeting synthetic CRISPR-Cas strain in *P. aeruginosa* produced no protection from phage lysis. Corroborating the importance of spacer choice (Xue et al., 2015). Experiments in *E. coli* showed that when the CRISPR-Cas system was directed against T5 lytic phage, only spacers that targeted pre-early genes (which enter the cell first upon infection to shut down cell activity (Davison, 2015)) resulted in effective CRISPR-Cas interference, and spacers targeting other regions provided no protection despite being effective at plasmid targeting (Strotskaya et al., 2017). This study also saw that for two of the lytic phages used – T4 and R1-37 – the CRISPR-Cas system in *E. coli* was unable to produce any protection against these phages (Strotskaya et al., 2017). This may have been due to DNA chemical modifications (which have been shown to inhibit CRISPR-Cas immunity (Vlot et al., 2018)) or some other unknown mechanism. In this study, two lysogenic phages JBD18 (fig S14) and JBD25 (fig S15) were both unable to be targeted by CRISPR-Cas interference despite the spacers being functional against plasmids. This may have been due to the spacer choice (as seen for the phage JBD63c) or due to another unknown mechanism. Given that JBD18 and JBD25 were described as ‘CRISPR sensitive’ in a previous study with the *P. aeruginosa* PA14 wildtype strain (Bondy-Denomy et al., 2013), the former spacer-based explanation is more likely. The synthetic CRISPR-Cas system also provided no protection against the jumbo phage phiKZ, which validates previous results that show phiKZ forms a proteinaceous shell around its DNA in the host which can prevent the action of type I CRISPR-Cas systems (Malone et al., 2020; Mozumdar et al., 2024).

The finding that CRISPR-Cas can only offer complete protection against lysis by some phages in low nutrient conditions highlights importance of considering the environment when testing phage infectivity and host range with defence systems. High nutrient, high growth environments are unlikely to reflect the ecological conditions of most bacteria-phage interactions. Other factors have also been shown to play a role in host infectivity, recently in *Acinetobacter baumannii* the host range of its phage Mystique was shown to vary between tests in liquid media and solid media (Alseth et al., 2024). Similarly in this study we found that the protective effects of CRISPR-Cas immunity produced more pronounced phenotypes in liquid rather than solid media experiments. Understanding these layered and nuanced interactions between phages and defence systems will allow us to better understand how bacteria may be able to evade and overcome the use of phages in agricultural or therapeutic settings. Larger screens of across more and diverse phages would facilitate searching for broad patterns and allow for the identification of predictive characteristics in phages that correlate with CRISPR-Cas immunity.

## Data availability

Data available upon request.

## Acknowledgements

We would like to thank Rosanna Wright and Michael Brockhurst for graciously sharing samples of the phages phiKZ, JBD88b, PA10P1, PA14P2, JBD10, JBD63c as well as the genomic sequences for PA10P1, PA14P2, JBD10, JBD63c phages that were not available on NCBI. This work was supported by the Biotechnology and Biological Sciences Research Council (sLoLa grant BB/X003051/1 to T.B.T. and E.R.W., and a South West Doctoral Training Programme BB/T008741/1 supporting J.F.E.), a UK Research and Innovation grant under the UK Government’s Horizon Europe funding guarantee (EP/X030377/1), a Royal Society Dorothy Hodgkin Research Fellowship (DHF\R\231005) to T.B.T. and the Philip Leverhulme Prize (PLP-2020-008) to E.R.W.

For the purpose of open access, the author has applied a ‘Creative Commons Attribution (CC BY) licence to any Author Accepted Manuscript version arising from this submission

## Author contributions

The synthetic CRISPR-Cas system was designed by JFKE and BNJW with oversight from ERW, TBT. All experiments were performed, and data analysed by JFKE. KC created annotated any unpublished phage genomes and performed phylogenetic analysis. The manuscript was written by JFKE with the methods and figure for the phylogenetic trees contributed by KC. All authors contributed to reviewing and editing the manuscript. Funding was provided by ERW and TBT.

## Supplementary methods Competition assays

Competition assays were performed with a 1:1 ratio of OD_600_ 1 normalised overnight cultures of *P. aeruginosa* in 10 ml LB broth at 0.1% (v/v). Competition cultures were incubated at 37 °C, shaking 24 hours. Bacterial titres were counted by spot-plating 20 µl of serially diluted culture on LB-agar plates with or without Gentamicin (10 μg/ml) in triplicates. The transformed strains only grew on LB-agar plates containing Gentamicin. The bacterial titre of the non-transformed strains was calculated as the difference in bacteria titre on the LB-agar plates and Gentamicin LB-agar plates, averaged across the technical repeats.

## Conjugation efficiency

The method was adapted from (Walker & Hatoum-Aslan, 2017) using a version of the pBBR1-MCS2 plasmid (Addgene plasmid # 85168 ; http://n2t.net/addgene:85168 ; RRID:Addgene_85168, modified version:) which was edited to swap the kanamycin resistance marker for tetracycline resistance (https://benchling.com/s/seq-HZj6BMhxcv6y6ta8UMTE). This plasmid was transformed into *E. coli* ST18 donor strain which requires 5-Aminolevulinic acid for growth. A version of the synthetic CRISPR-Cas system was created which had a spacer targeting against a short sequence adjacent to the multiple cloning site (table S2), and then transformed into *P. aeruginosa*. Strains were incubated and grown in 10 ml LB for 16 hours before being normalised to an OD_600_ of 0.4 in LB. 400 μl of each recipient strain and donor strain were combined, spun at 8000 rpm for 3 minutes and resuspended in 1 ml LB, spun again and resuspended in 50 μl and plated on a cellulose filter on 5-Aminolevulinic acid (50 μg/ml) LB-agar plates and grown overnight to allow conjugation. The following day each filter was vortexed in 3 ml LB to release bacteria. The resuspended bacterial culture was diluted in LB and 10 μl spots were plated in technical triplicate on both VBMM-agar plates (to count total *P. aeruginosa* bacterial titre) and tetracycline (15 μg/ml) VBMM-agar plates (to determine the titre of successful transconjugants). VBMM-agar plates (Choi & Schweizer, 2006) prevent *E. coli* growth, thus selecting for *P. aeruginosa* growth only. Conjugation efficiency was calculated as the number of successful transconjugants divided by the total number of bacteria.

The same method was used to validate the efficiency of interference of each phage targeting spacer 1 in the synthetic CRISPR-Cas system. pBBR1-MCS2 tetracycline modified plasmids were edited to include a 100-300 base pair fragment around the protospacer in the phage genome (table S3) via PCR then restriction-ligation using the SacI and HindIII recognition sites in the plasmid multiple cloning site. Plasmids were then validated via PCR and sequencing (tet-pBBR1-MCS2 F & R table S2).

## Efficiency of plating

Strains with a phage targeting spacer, non-targeting spacer, the Tn7 insertion elements with no cargo genes, and the CRISPR-Cas knockout *P. aeruginosa* strains were plated as a lawn (100 μl of overnight culture in 4 ml 0.4% LB-agar plated on 1.5% LB-agar plates) upon which phage dilutions were spotted. The number of plaques were counted the following day from which efficiency of plating was calculated. Efficiency of plating is defined as the titre of phage on each strain / average phage titre plated on the control strain (CRISPR-Cas knock-out). The method was repeated for each phage in the panel of 13 phages using the CRISPR-Cas knockout, non-targeting spacer strains as well as a strain with two spacers matching each phage (table S3).

## Evolution with DMS3vir to determine spacer acquisition

A three-day evolution experiment was performed to test if spacer acquisition could occur with the *P. aeruginosa* synthetic CRISPR-Cas system and phage DMS3vir. The CRISPR-Cas knockout strain as well as strains with either a non-targeting, phage targeting, or phage primed spacers were inoculated into glass vials with 6 ml M9 media in six independent replicates. These replicates were transferred 1:100 into fresh M9 media the next day and infected with 10^4^ plaque forming units (pfu) per ml. Cultures were then incubated at 37 °C, shaking at 200 r.p.m overnight. During the three-day experiment, cultures were transferred 1:100 into fresh medium every 24 hours. The type of resistance mechanism that had evolved was determined at three days post infection by randomly picking twelve colonies for each of the six biological replicates for each strain, growing them overnight in 150 μl LB then streaking the cultures through DMS3*vir* and DMS3*vir-acr*IF1 phage. Surface modification was confirmed by broad-range resistance to both phages. CRISPR-Cas immunity was confirmed by resistance to DMS3*vir* but sensitivity to DMS3*vir-acr*IF1. From these immune samples, spacer acquisition was detected by PCR using the primers Tn7 F and CRISPR array R (table S2) followed by gel electrophoresis in a 2% agarose gel to resolve DNA size differences (a subset of which were confirmed by Sanger sequencing).

## Phage phylogeny

Complete phage genomes were retrieved from NCBI on 16/04/2024 by filtering for viruses and searching “Pseudomonas aeruginosa”, “phage”, and “complete”. In total, 1014 phage genomes were retrieved, and filtered down to 854 based on whether their metadata confirmed that they were isolated from *P. aeruginosa*. All phage genome sequences were annotated with Pharokka (v1.7.1) (Bouras et al., 2023). The amino acid sequences of the terminase large subunit (TerL) and the major capsid protein were chosen as these were present in 721 and 639 phages, respectively. The amino acid sequences were aligned using MUSCLE (Edgar, 2004), and phylogenetic trees were constructed using the maximum likelihood algorithm using MEGA11 (Tamura et al., 2021). Trees were visualised using Microreact (Argimón et al., 2016).

## Phage strains used in study

**Table S1.** Panel of phages the synthetic CRISPR-Cas system was screened against

| Phage | NCBI reference |
| --- | --- |
| DMS3 | NC_008717.1 (Zegans et al., 2009) |
| JBD5 | NC_020202.1 (Bondy-Denomy et al., 2013) |
| JBD10 | Shared by Brockhurst lab (Bondy-Denomy et al., 2016) |
| JBD88b | KM389212.1 (Bondy-Denomy et al., 2016) |
| PA10P1 | Shared by Brockhurst lab (Wright et al., 2018) |
| PA14P2 | Shared by Brockhurst lab (Wright et al., 2018) |
| LMA2 | NC_011166.1 (P. Ceyssens et al., 2009) |
| JBD63c | Shared by Brockhurst lab (Bondy-Denomy et al., 2016) |
| LBP1 | NC_027298.1 (Lavigne lab) |
| JBD18 | NC_027986.1 (Cady et al., 2012) |
| JBD25 | NC_027992.1 (Cady et al., 2012) |
| PhiKZ | NC_004629.1 (Krylov et al., 2021) |

## Primers and sequences used

**Table S2.**
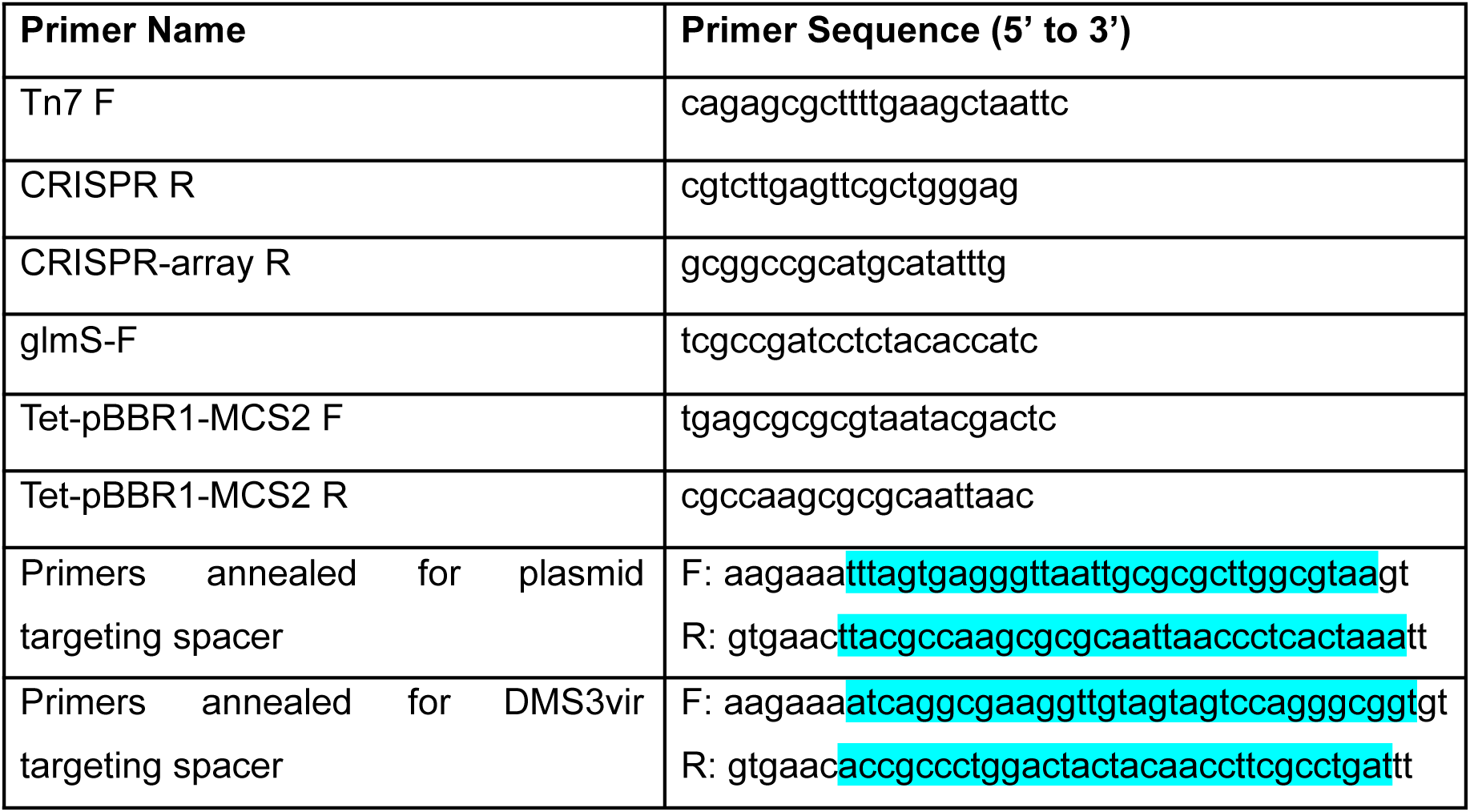
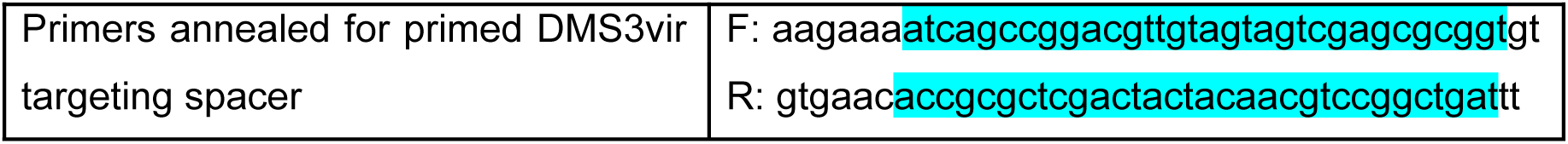
Primer table listing the sequences of primers used throughout the study to validate cloning and create spacer fragments.

**Table S3.**
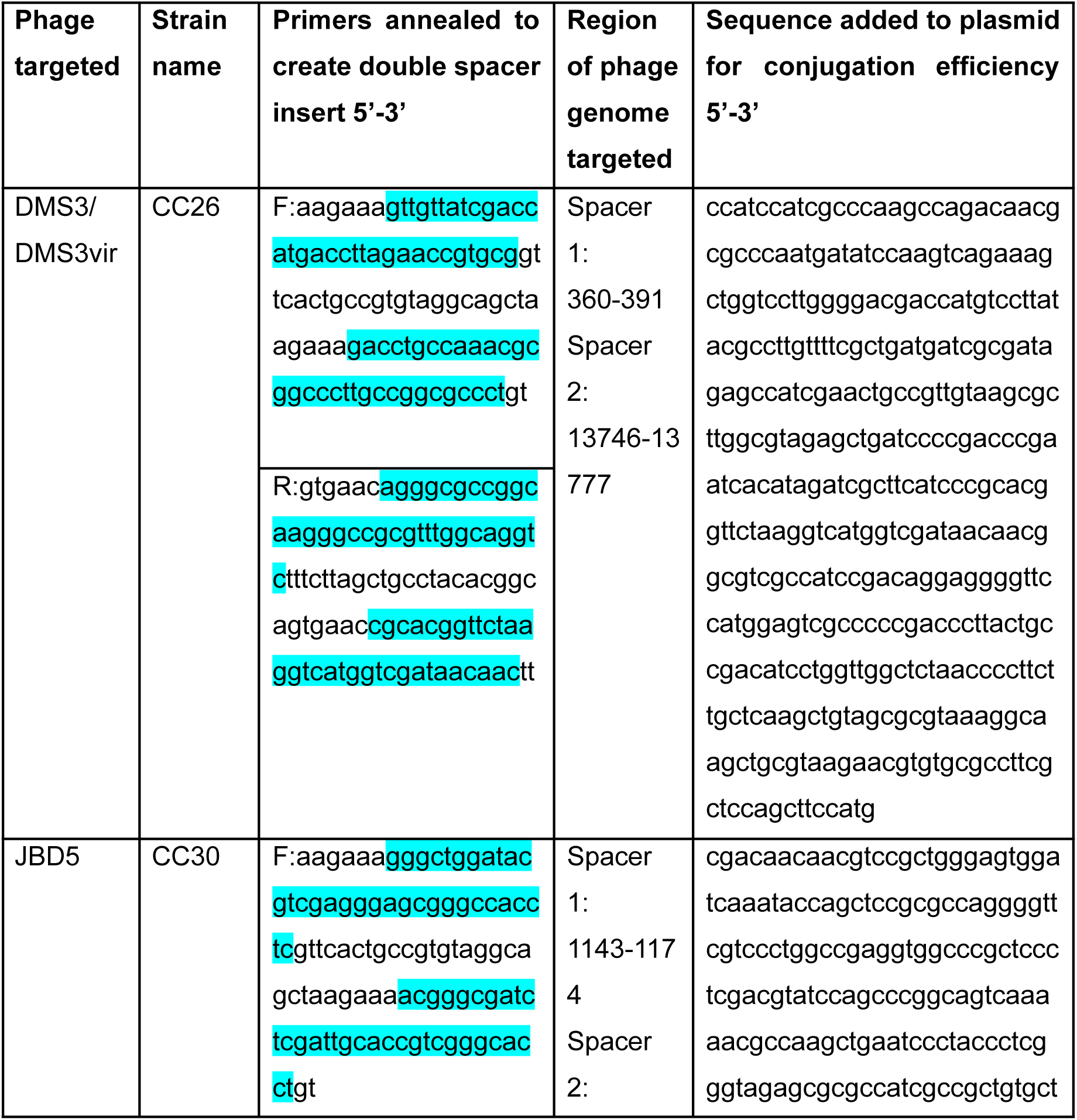
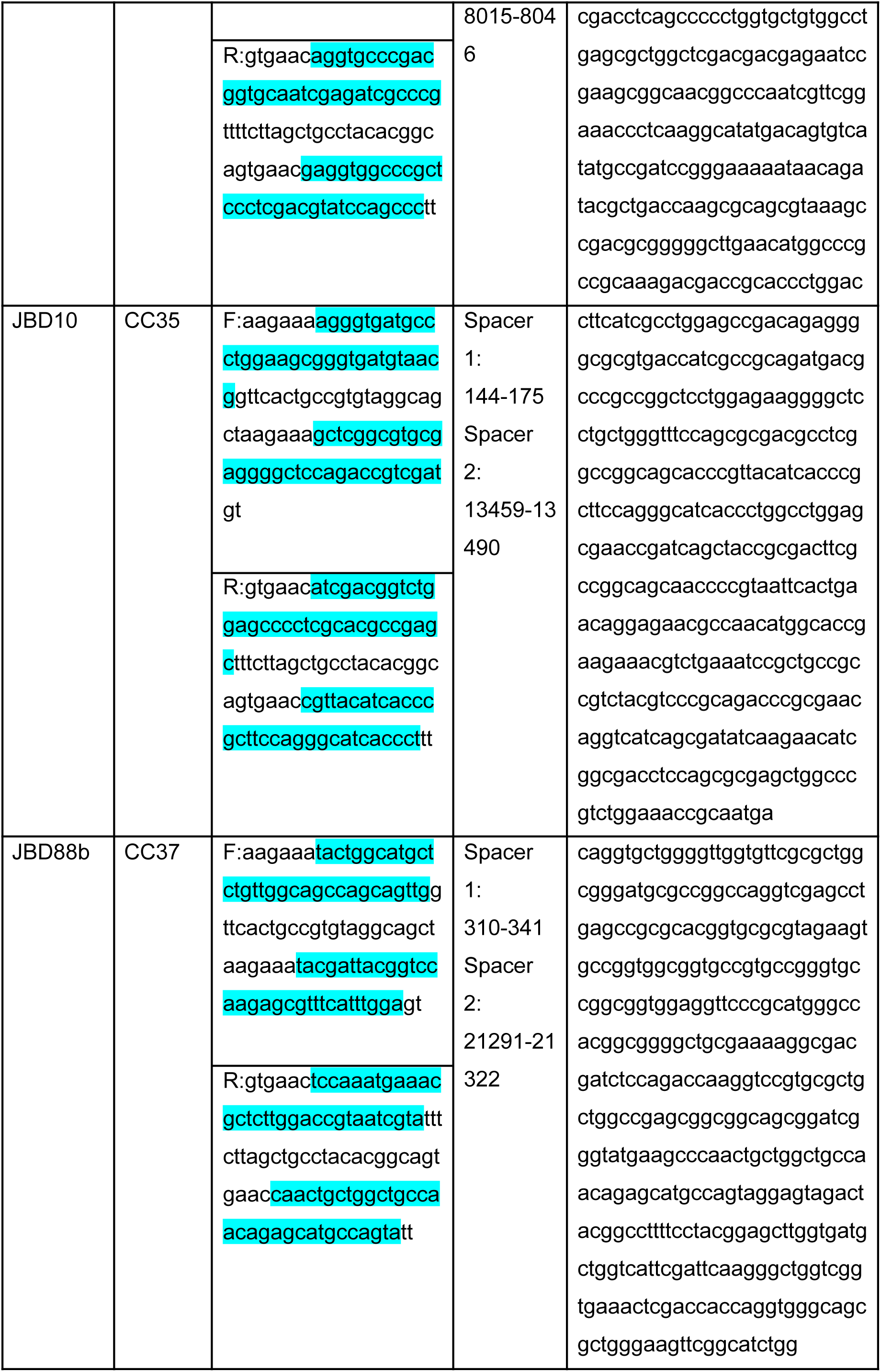
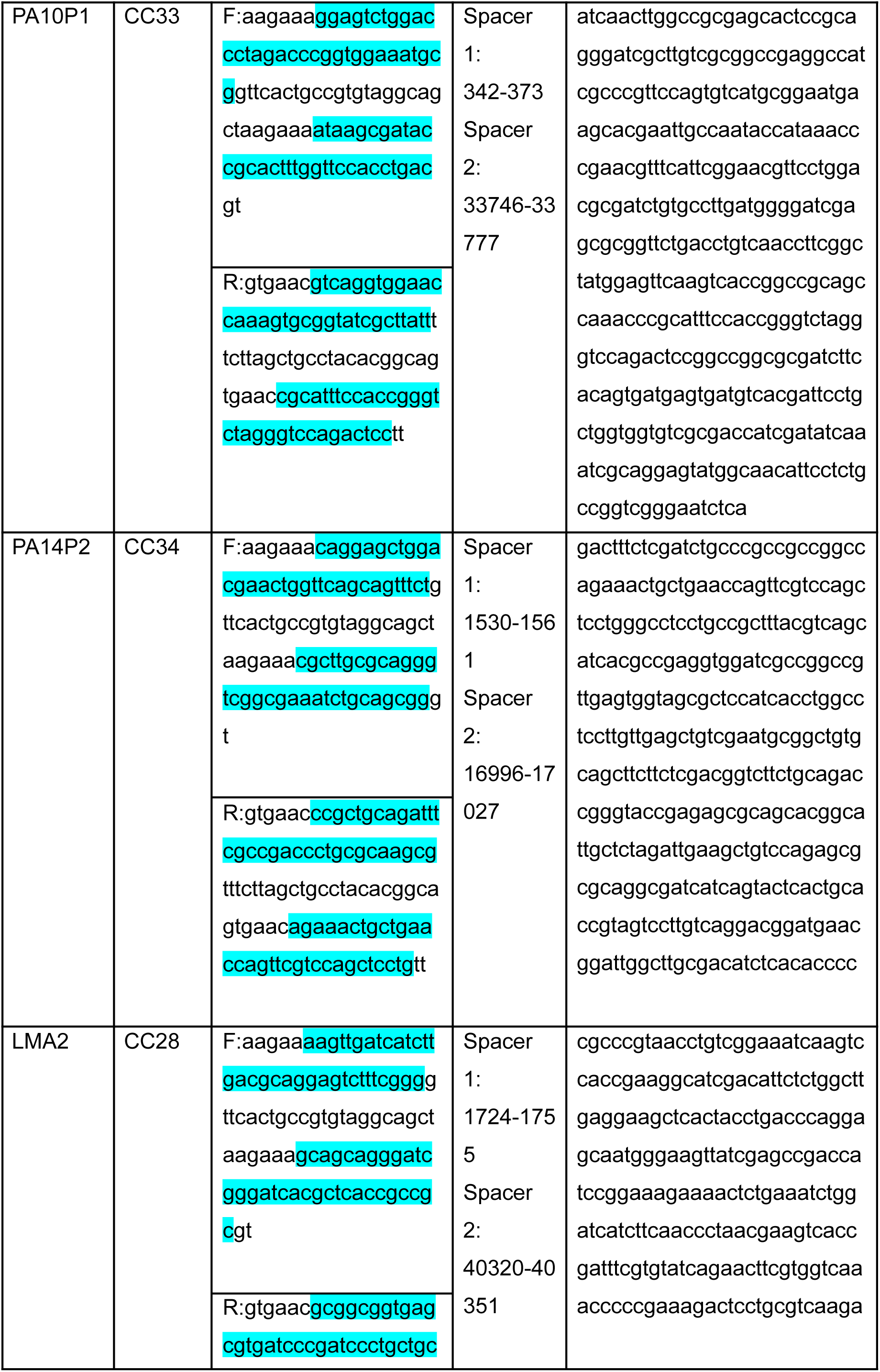
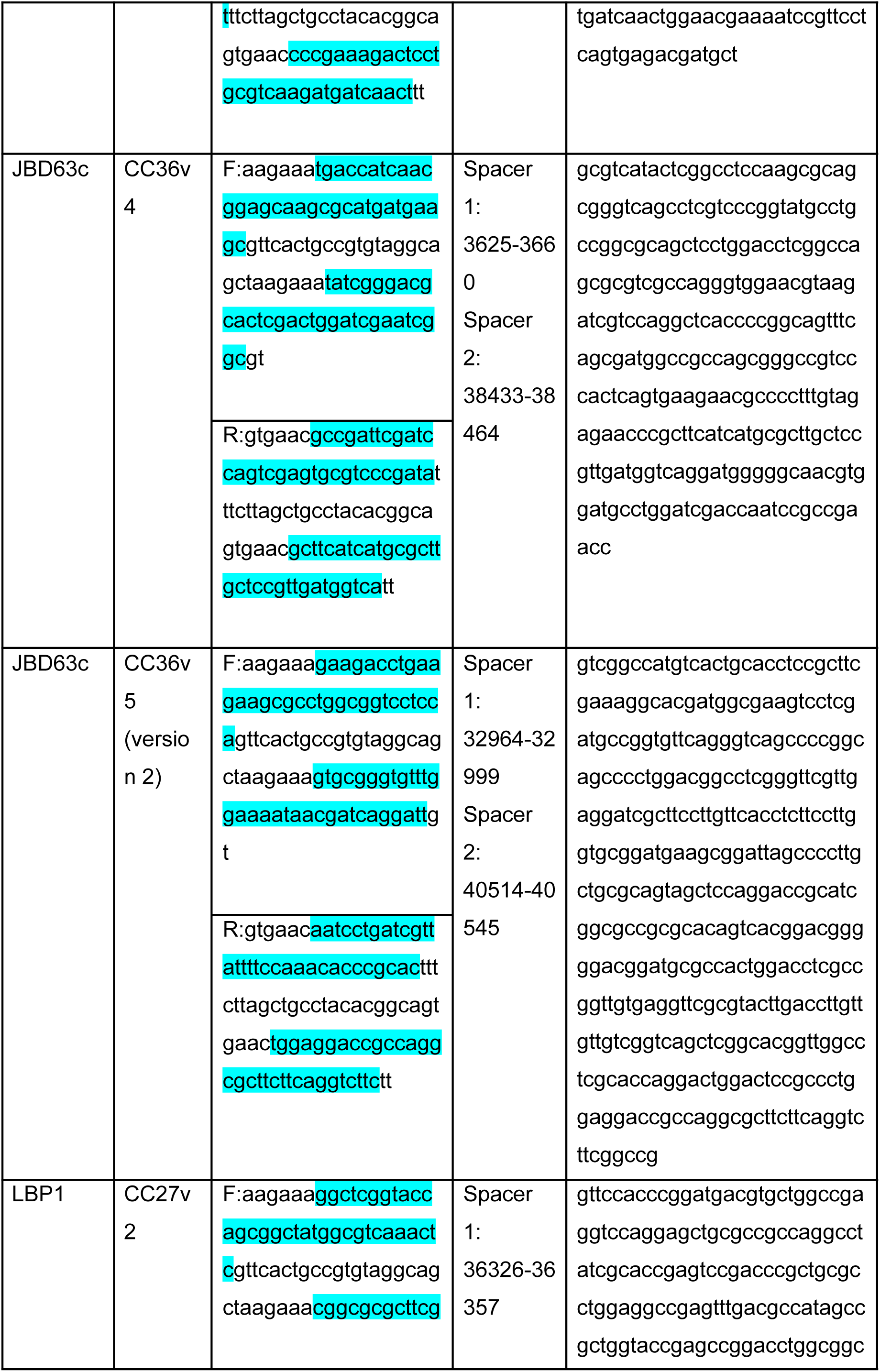
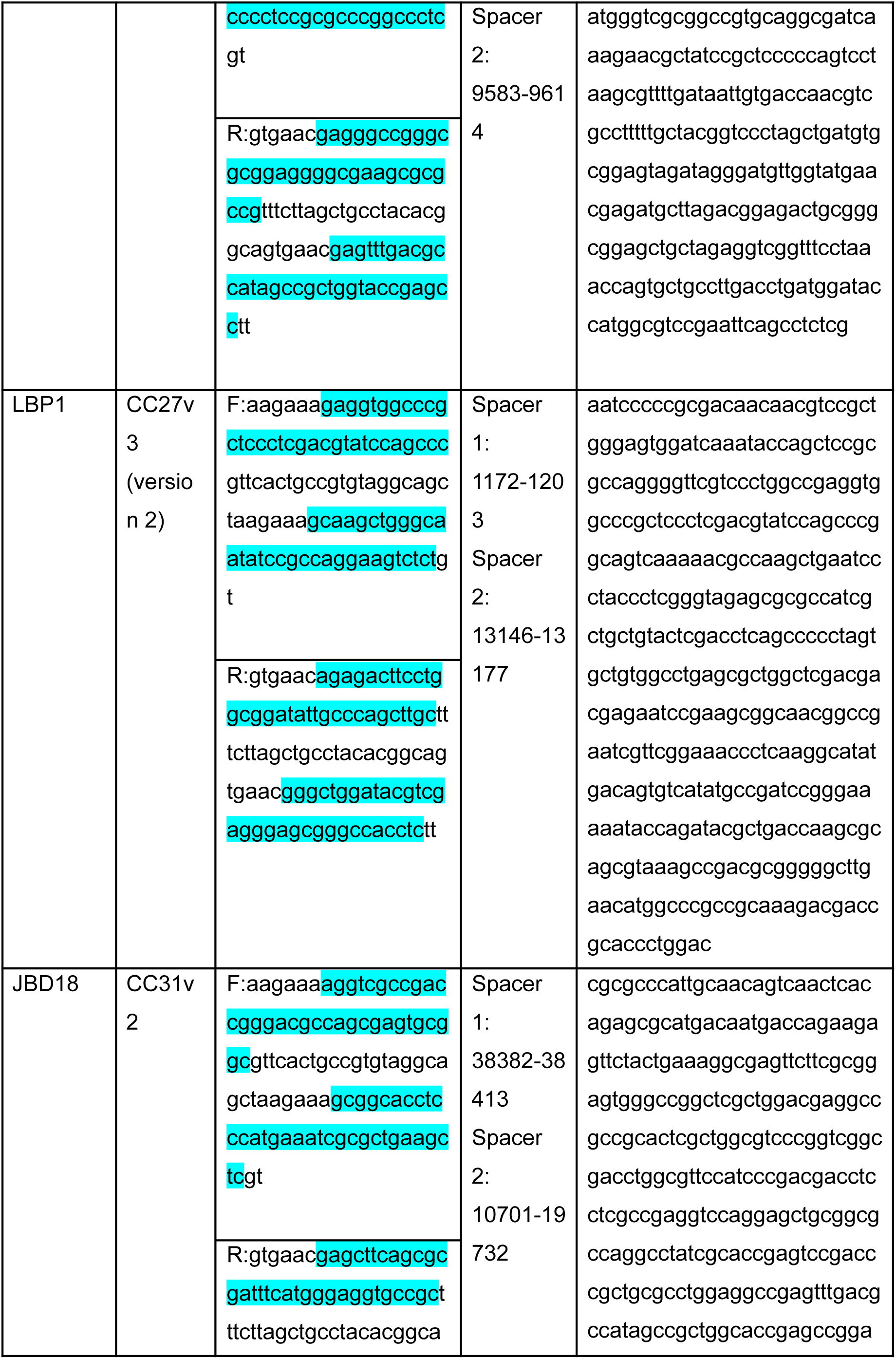
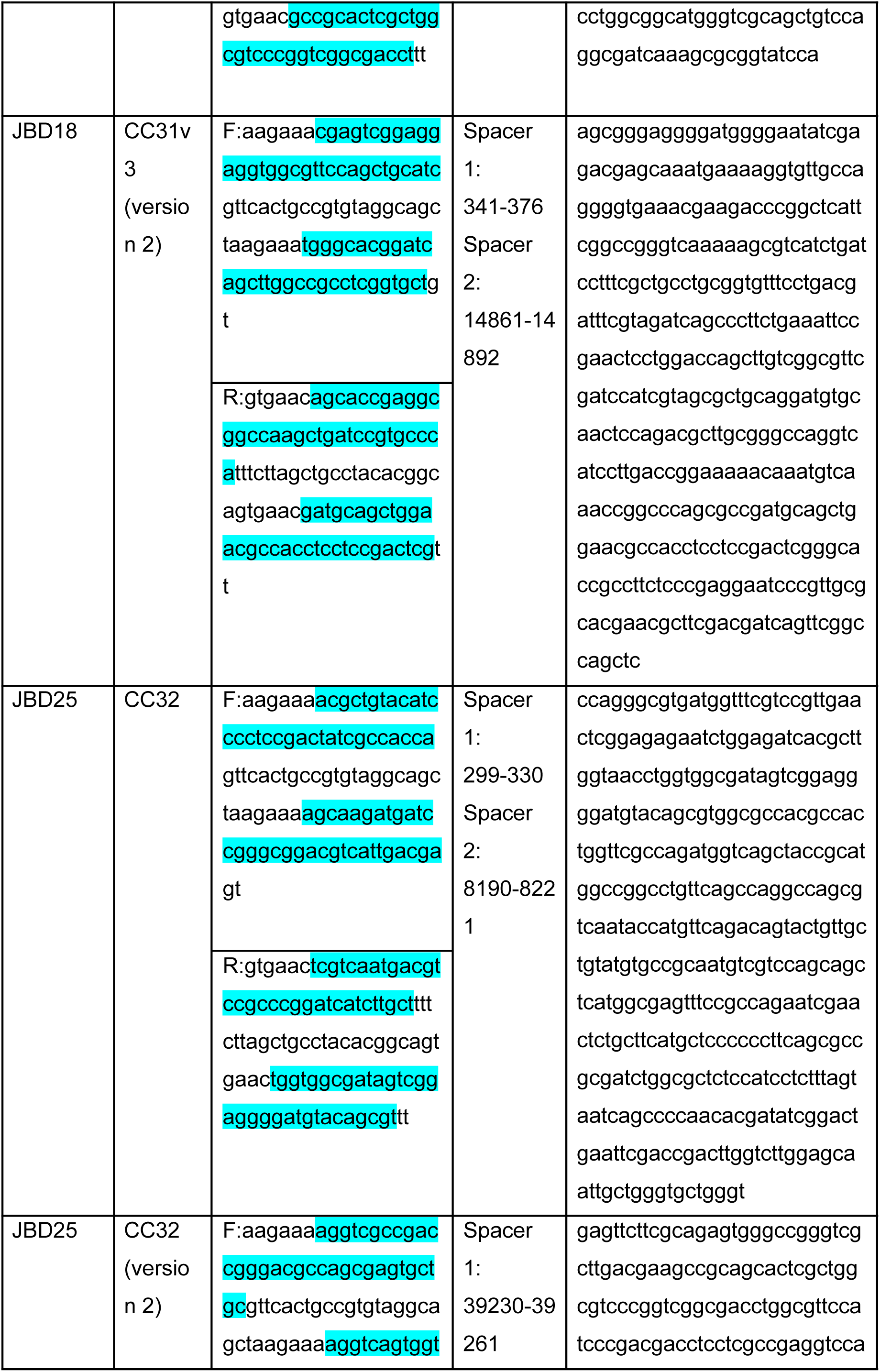
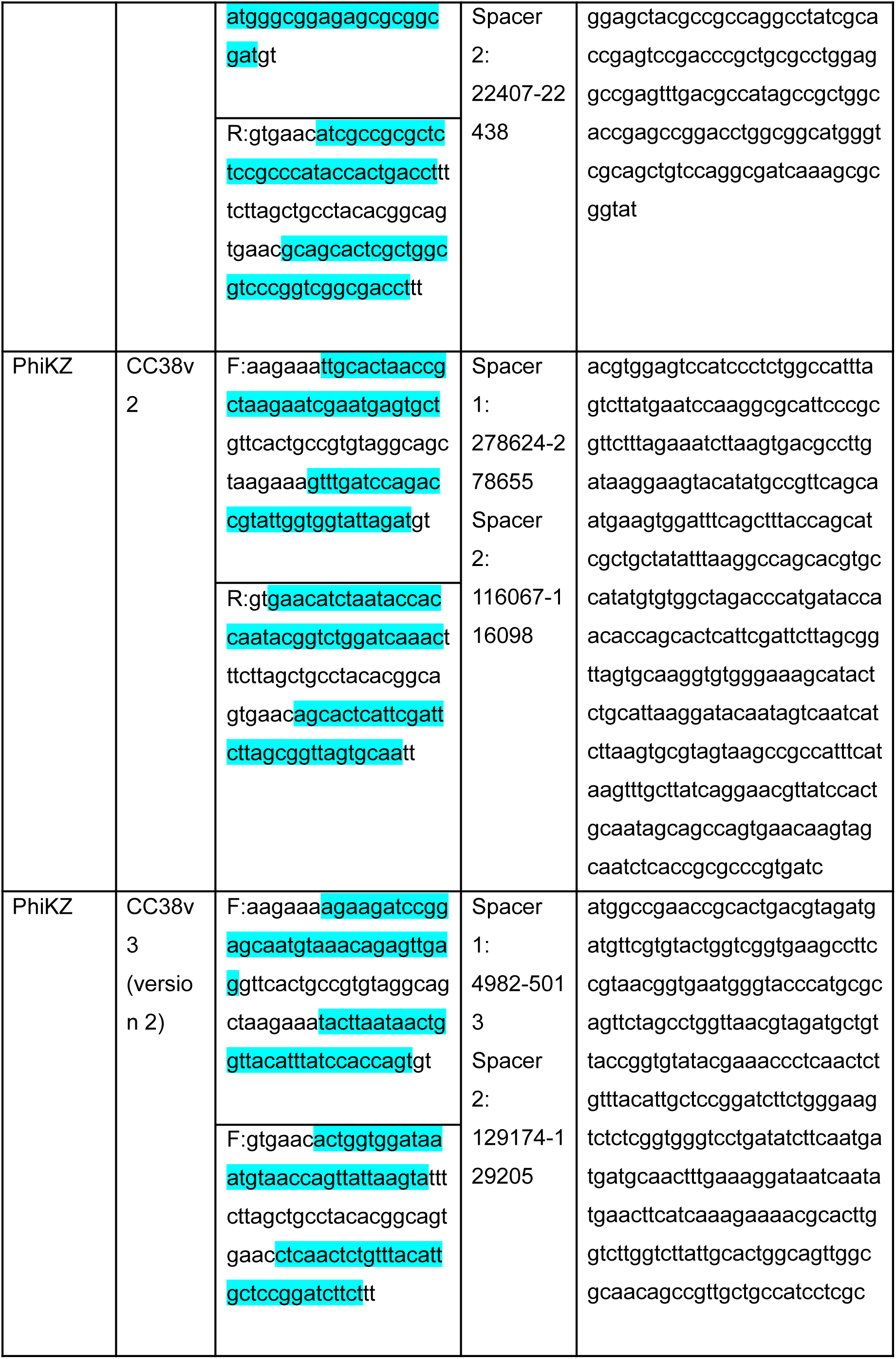
Primers annealed to create synthetic CRISPR-Cas systems that target the various phages used. The two spacer sequences in the primers annealed to clone each CRISPR array are highlighted in blue. Spacers were designed to target different regions in the genome and regions containing spacer 1 for each strain were cloned into plasmid for further validation via conjugation efficiency. These were ligated into the plasmid pUC18T-mini-Tn7T-Gm-CC01 https://benchling.com/s/seq-Z1606qDdCNDkh3iX1Kl5?m=slm-7kmTa3tzY5ehw9IuZ0AN which contained the ‘non-targeting’ spacer 5’ gtcttcagttcacgcgtagctcaaagaagac 3’.

**Figure S1.**
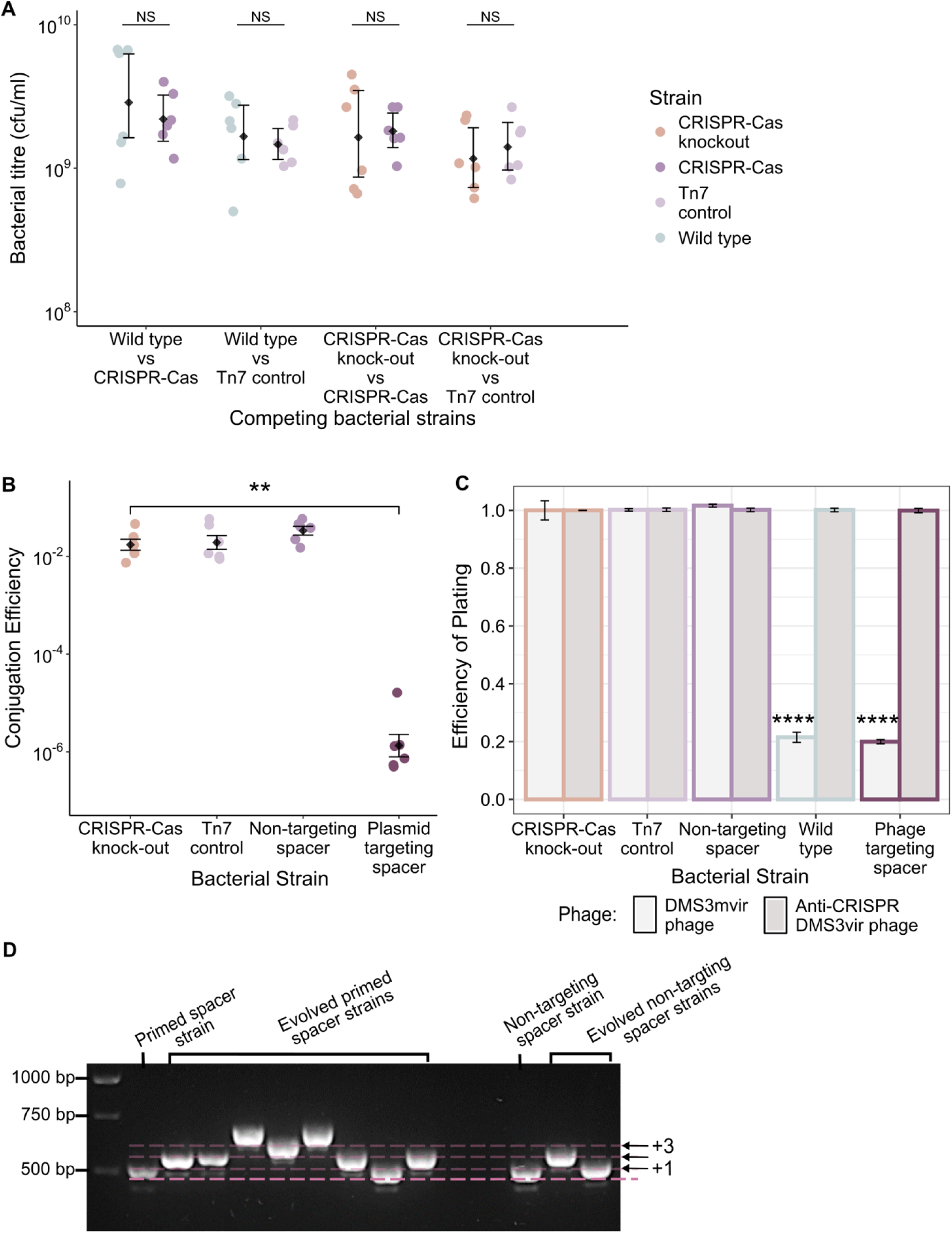
– **Validation of the synthetic CRISPR-Cas system** (A) Competition assay to assess the potential fitness costs due to the addition of the synthetic CRISPR-Cas system via the mini-Tn7 integration system. Strains were mixed in a 1:1 ratio and competed over 24 hours. The gentamicin resistance gene in the mini-Tn7 integration system allowed for selection during plating. The resultant bacterial titre (cfu/ml) of each strain plotted for each competition experiment. The strains competed are denoted by the colour. Mean values are given (N=6) and error bars represent 95% confidence intervals. Statistical significance between the two strains in each competition experiment is denotes by ‘ns’ for non-significant. Dots show individual data points for each replicate. A paired sample t-test (using a Benjamini-Hochberg adjustment of p-values) was used to assess if any of the strains grew to a higher bacterial titre during each competition experiment. There was no significant difference in bacterial titre between each pair of strains in each competition experiment. Therefore, this indicates that cloning in the synthetic CRISPR-Cas system via the mini-Tn7 integration system doesn’t not carry a constitutive fitness cost. The wildtype and CRISPR-Cas knock-out strains could not be competed due to the lack of a suitable selection marker between the two strains. (B) Validation of the ability of the synthetic CRISPR-Cas system to target foreign DNA for cleavage using conjugation efficiency of plasmids. Conjugation efficiency experiment calculated as the number of successful transconjugants with the tetracycline resistant pBBR1-MCS2 plasmid. Black diamonds denote mean values (N=6), and error bars represent standard error around the mean. Individual data points are shown in dots coloured by strain. The difference in the plasmid conjugation efficiency between each strain was assessed by ANOVA analysis (F_3,20_ = 191.3, p < 6.841e-15, Adjusted R^2^ = 0.9613, data was log_10_ transformed). T-test comparing conjugation efficiency between the CRISPR-Cas knock-out and plasmid targeting spacer showed a significant decrease in conjugation efficiency (** denotes p<0.01), with the number of transformed cells dropping 10,000-fold. Whereas the Tn7 control (that contains the Tn7 integration elements but no cargo genes) and non-targeting spacer strain all conjugated plasmids with similar efficiency to the CRISPR-Cas knock-out strain. (C) Validation of the ability of the synthetic CRISPR-Cas system to act as a phage defence system. Efficiency of plating is plotted for each strain against both DMS3mvir-AcrIE3 and DMS3vir-AcrIF1 phage. The DMS3mvir-AcrIE3 phage matches the sequence of the first spacer of the native CRISPR 2 array in wildtype *P. aeruginosa*, the same spacer was engineered into the phage-targeting spacer synthetic CRISPR-Cas system. The bar chart is shaded by phage type and outlined in colours corresponding to the strain type. The mean phage titre on the CRISPR-Cas knock-out strain was used as the reference when calculating efficiency of plating. Mean values (N=3) are plotted and error bars represent 95% confidence intervals. The difference in efficiency of plating for each strain on DMS3mvirAcrIE3 was assessed by ANOVA analysis (F_4,10_ = 2459, p < 6.485e-15, Adjusted R^2^ = 0.9986). Tukey pairwise comparison testing comparing each strain to the CRISPR-Cas knock-out reference strain, showed that the Tn7 control and non-targeting spacer containing strain of *P. aeruginosa* efficiency of plating was not significantly different. However, both the wild type *P. aeruginosa* and the strain containing a synthetic CRISPR-Cas with a phage targeting spacer had significantly reduced efficiency of plating (p < 0.0001, denoted by ‘****’). Phage levels dropped from titres of 10^10^ pfu/ml to 10^2^ pfu/ml when plated on these two strains. This shows that the synthetic CRISPR-Cas can perform comparably to the native counterpart. When this experiment was repeated with a version of the DMS3vir phage carrying an anti-CRISPR gene (DMS3mvir-acrIF1) there was no significant difference in efficiency of plating between all strains, demonstrating that the previously observed phage defence was due to the CRISPR-Cas system rather than another mechanism. (D) Validation of the ability of the synthetic CRISPR-Cas system to acquire new spacers. The CRISPR-Cas knock-out, synthetic CRISPR-Cas with non-targeting, phage targeting, and phage primed spacer strains were co-evolved with the DMS3vir phage in M9 media for three days, to ensure sufficient time for spacer acquisition to occur and undergo selection. The experiment was performed in nutrient limited media as this has previously been shown to favour the evolution of CRISPR-Cas immunity in *P. aeruginosa* PA14 strains (Westra et al., 2015). DMS3vir phages were added at the beginning of the experiment (day 0) at a titre of ∼10^4^ pfu/ml. The resistance mechanisms were then profiled at three days post infection using twelve randomly sampled individual colonies for each strain. Colonies that were resistant when streaked through DMS3vir phage but not DMS3vir-acrIF1 phage were inferred to be protected by CRISPR immunity. The agarose gel pictured shows changes in DNA band size due to spacer acquisition relative to ancestral *P. aeruginosa* PA14 strains after PCR of a subset of the immune clones. The lowest dashed line aligns to the size of the amplified CRISPR-array with a single spacer (524 base pairs) with the above dashed lines at each 60 base pair increment that indicate the incorporation of 1, 2, or 3 new spacers into the CRISPR array. Clones with expanded CRISPR arrays in the primed spacer strain indicated primed spacer acquisition, and the single non-targeting spacer clone with an expanded array demonstrates naïve spacer acquisition.

**Figure S2.**
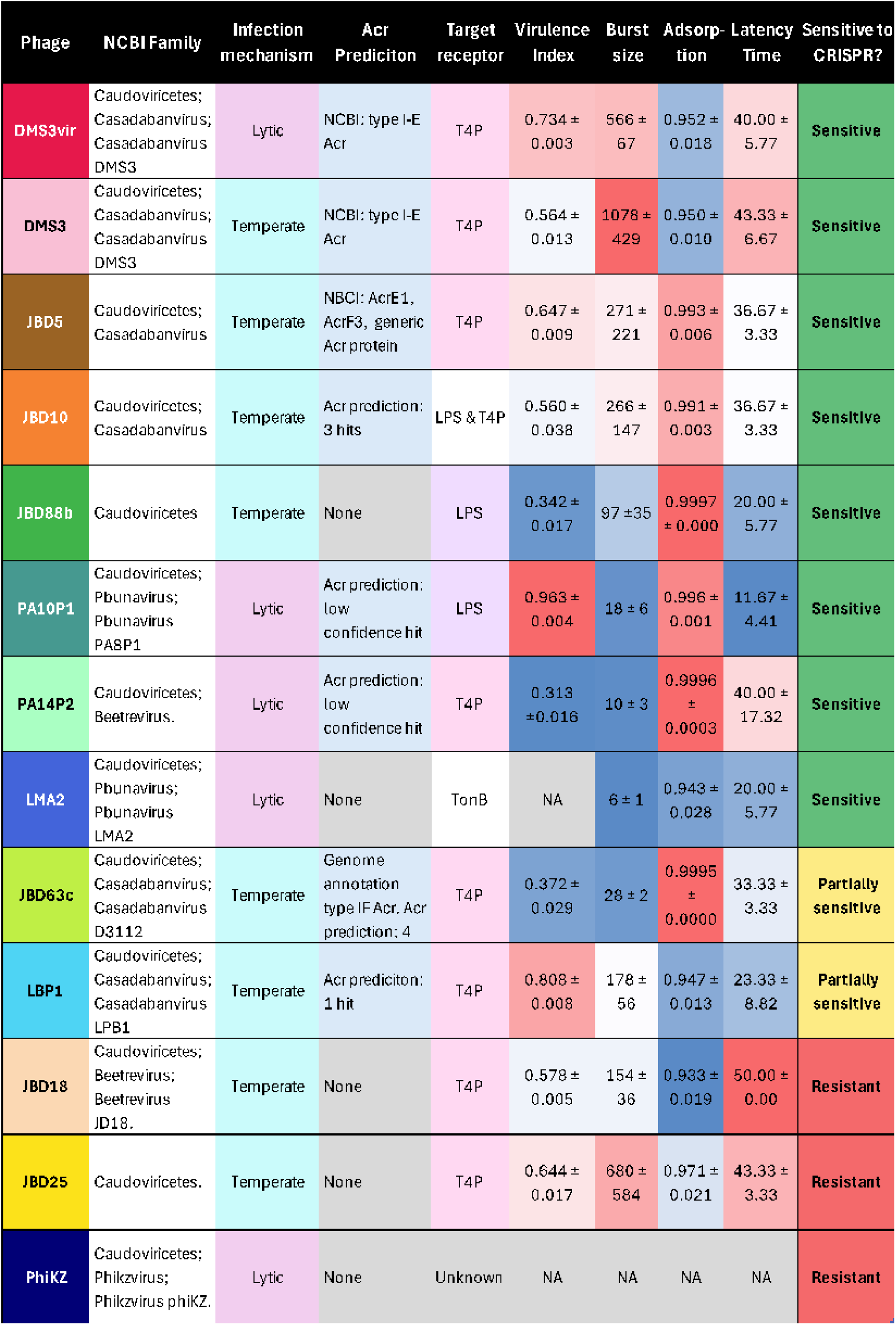
– **There is no clear predictive phage variable that correlates to the ability of CRISPR-Cas to provide protection through interference.** Table comparing phage characteristics and the susceptibility of each phage to interference by the synthetic CRISPR-Cas system. Phage names are coloured by type/species using the same colour scheme used in figure 3 and 4 as well as in figure S4-15A. Phage species identification are given from the last taxa common to all species (Caudoviricetes) as listed on their NCBI reference. The infection mechanism is coloured by type – lytic or temperate. Whether any anti-CRISPR genes are predicted in the genome is listed using first, any labelling of anti-CRISPR genes on NCBI, then any anti-CRISPR genes labelled using Pharokka, and then if any anti-CRISPR genes were predicted by AcrFinder (Yi et al., 2020) and the AntiDefence Finder function on DefenseFinder (Tesson et al., 2022, 2024). The receptor that each phage binds to on the cell surface is listed, T4P – type IV pilus, LPS – lipopolysaccharide membrane, TonB – TonB-dependent receptor. For T4P binding phages this could be validated by lack of infection on type IV pilus knockout strains. Virulence index (Storms et al., 2020) was calculated from virulence curves in figure 3D, mean values are given ± standard error and coloured by value with red for highest virulence and blue for lowest. Burst size (phages released per cell) is summarised from figure 3A given as mean values ± standard error and coloured by value with red for highest burst size and blue for lowest. Adsorption (proportion of phages adsorbed) is summarised from figure 3B and given as mean values ± standard error and coloured by value with red for highest adsorption values and blue for lowest. Latency time (time to burst) is summarised from figure 3C and given as mean values ± standard error and coloured by value with red for longest latency time and blue for shortest. Whether or not the phage is sensitive to interference by the synthetic CRISPR-Cas system with spacer targeting the phage is shown with green for complete CRISPR-Cas immunity, yellow if immunity is affected by environment or spacer choice, and red if no CRISPR-Cas immunity was observed in this study.

**Figure S3.**
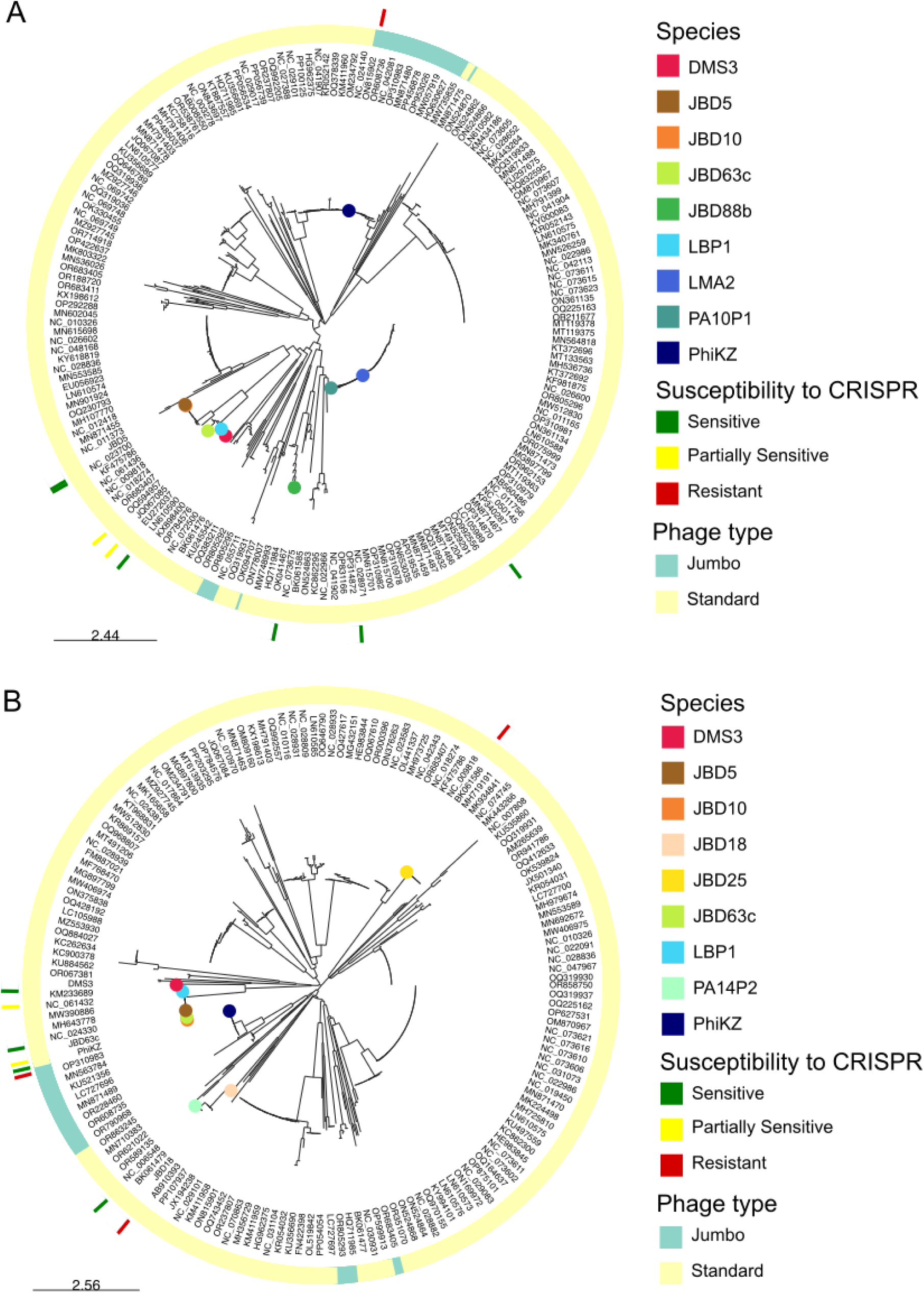
– **Phylogenetic trees of Pseudomonas aeruginosa-infecting phages** based on the amino acid sequence of the terminase large subunit (A) and the major capsid protein (B). The end branch for each phage in the panel is indicated with a circle corresponding in colour to the phages. The phage type for each terminal taxa, either jumbo (>200kb) or standard genome size, are depicted in teal or yellow respectively. Whether further testing showed that the phages were susceptible to interference by the synthetic CRISPR-Cas system (sensitive – green), susceptible to interference under certain conditions (partially sensitive – yellow), or not susceptible to interference by the CRISPR-Cas system (resistant-red) is depicted by a block on the outermost circle corresponding to the terminal taxa for each phage in the phylogenetic tree.

**Figure S4.**
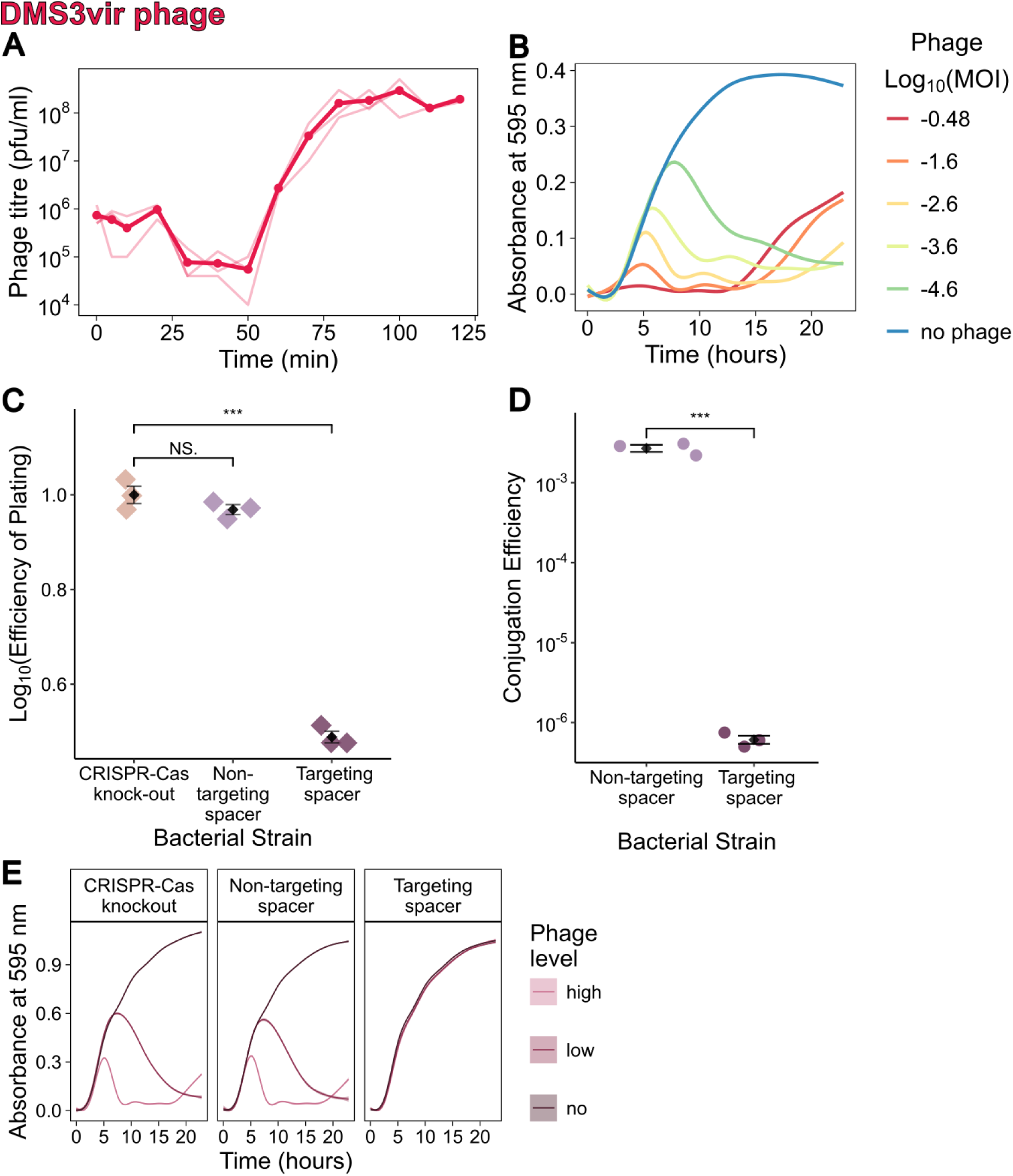
– **DMS3vir phage characteristics**. (A) One-step growth curve. Assays were performed at a starting MOI of approximately 0.1. Samples were treated with chloroform to lyse cells, meaning both free phages and mature phages inside cells were measured. The mean (N=3) dots are joined by a bold line with individual samples shown by thin lines. Phages were plated on *P. aeruginosa* wildtype cultures to calculate phage titre (pfu/ml) at each time point. (B) Growth curves in high nutrient liquid media at a range of phage MOIs to calculate virulence with *P. aeruginosa* wildtype with *P. aeruginosa* wildtype. Smoothened mean (N=3) absorbance at OD_595_ lines are coloured rainbow scale red to blue by decreasing log transformed MOI across over 23 hours. (C) Efficiency of plating experiment to test interference by the synthetic CRISPR-Cas system on solid media. Black diamonds denote the mean (N=3), with individual replicate denoted by diamonds coloured by strain type. Error bars give the standard error interval. The CRISPR-Cas knock-out strain is used as the reference strain when calculating efficiency of plating. Statistical testing are pairwise t-tests (NS. non-significant, *** p<0.001). On the non-targeting spacer strain the phages grew similarly to the CRISPR-Cas knock-out strain, whereas the strain with a targeting spacer led to a significant decrease in plated phage titre. (D) Conjugation efficiency experiment to validate the phage targeting spacer. The region of the phage genome containing the protospacer targeted by spacer 1 in the synthetic CRISPR array was incorporated into a plasmid. Black diamonds denote the mean (N=3), with individual replicate denoted by dots coloured by strain type. Error bars give the standard error interval. Compared to the non-targeting spacer strain, there was a significant decreased in the proportion of the cell population that took up the protospacer-containing plasmid for the targeting spacer strain (t -test, *** p<0.001). (E) Growth curves for each strain – CRISPR-Cas knock-out, non-targeting spacer and targeting spacer strains in the presence of no phage, a low level of phage (10^4^ pfu/ml), or a higher level of phage (10^6^ pfu/ml). Smoothened mean (N=6) OD_595_ absorbance lines were plotted, coloured by phage level, over a 23-hour growth cycle in nutrient rich liquid media. Each line has ± standard error shaded around the mean line. When *P. aeruginosa* possess the synthetic CRISPR-Cas system with a phage targeting spacer similar growth dynamics are seen with and without phages present.

**Figure S5.**
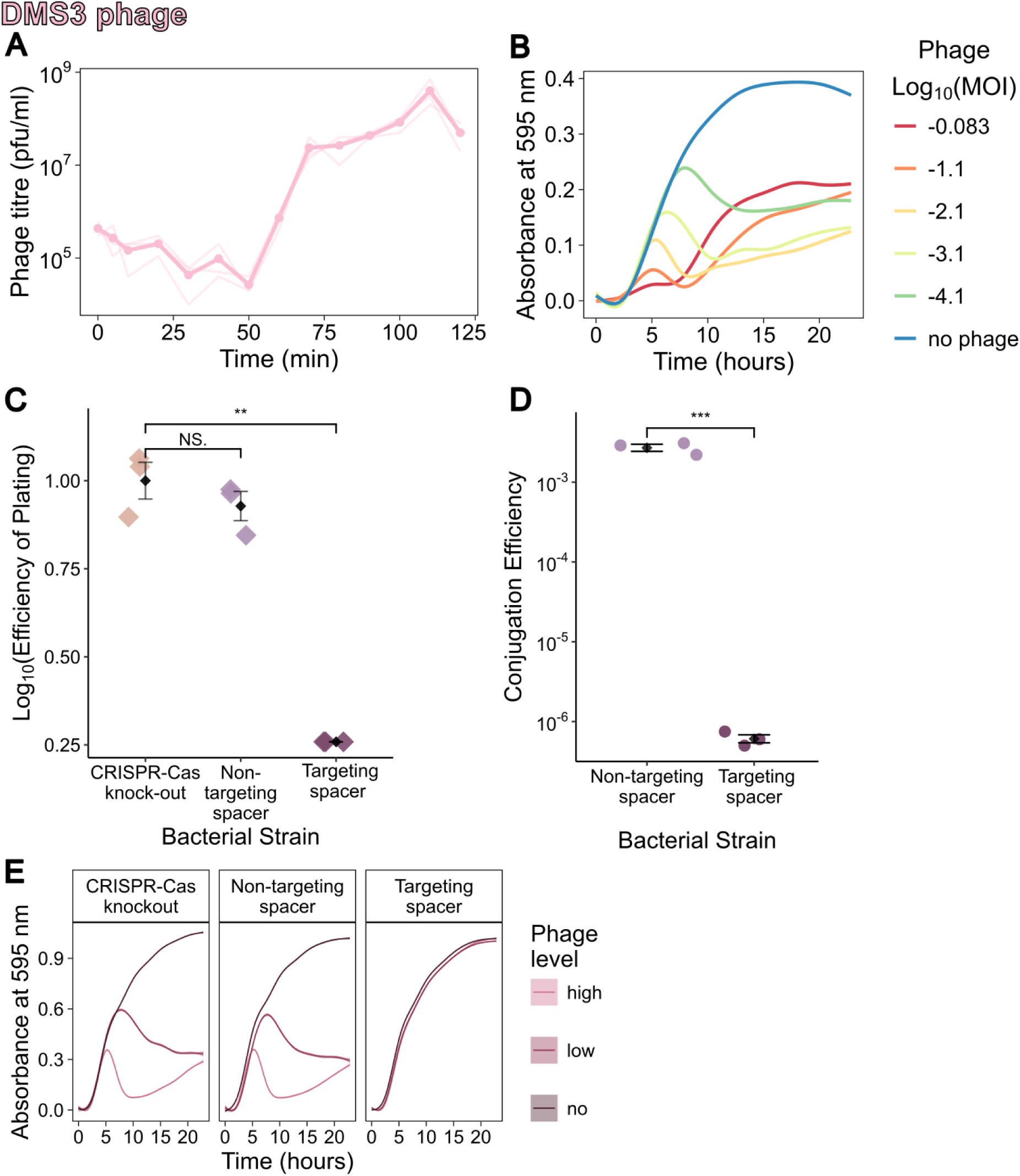
– **DMS3 phage characteristics**. (A) One-step growth curve. Assays were performed at a starting MOI of approximately 0.1. Samples were treated with chloroform to lyse cells, meaning both free phages and mature phages inside cells were measured. The mean (N=3) dots are joined by a bold line with individual samples shown by thin lines. Phages were plated on *P. aeruginosa* wildtype cultures to calculate phage titre (pfu/ml) at each time point. (B) Growth curves in high nutrient liquid media at a range of phage MOIs to calculate virulence with *P. aeruginosa* wildtype. Smoothened mean (N=3) absorbance at OD_595_ lines are coloured rainbow scale red to blue by decreasing log transformed MOI across over 23 hours. (C) Efficiency of plating experiment to test interference by the synthetic CRISPR-Cas system on solid media. Black diamonds denote the mean (N=3), with individual replicate denoted by diamonds coloured by strain type. Error bars give the standard error interval. The CRISPR-Cas knock-out strain is used as the reference strain when calculating efficiency of plating. Statistical testing are pairwise t-tests (NS. non-significant, ** p<0.01). On the non-targeting spacer strain the phage grew similarly to the CRISPR-Cas knock-out strain, whereas the strain with a targeting spacer led to a significant decrease in plated phage titre. (D) Conjugation efficiency experiment to validate the phage targeting spacer. The region of the phage genome containing the protospacer targeted by spacer 1 in the synthetic CRISPR array was incorporated into a plasmid. Black diamonds denote the mean (N=3), with individual replicate denoted by dots coloured by strain type. Error bars give the standard error interval. Compared to the non-targeting spacer strain, there was a significant decreased in the proportion of the cell population that took up the protospacer-containing plasmid for the targeting spacer strain (t -test, *** p<0.001). (E) Growth curves for each strain – CRISPR-Cas knock-out, non-targeting spacer and targeting spacer strains in the presence of no phage, a low level of phage (10^4^ pfu/ml), or a higher level of phage (10^6^ pfu/ml). Smoothened mean (N=6) OD_595_ absorbance lines were plotted, coloured by phage level, over a 23-hour growth cycle in nutrient rich liquid media. Each line has ± standard error shaded around the mean line. When *P. aeruginosa* possess the synthetic CRISPR-Cas system with a phage targeting spacer similar growth dynamics are seen with and without phages present.

**Figure S6.**
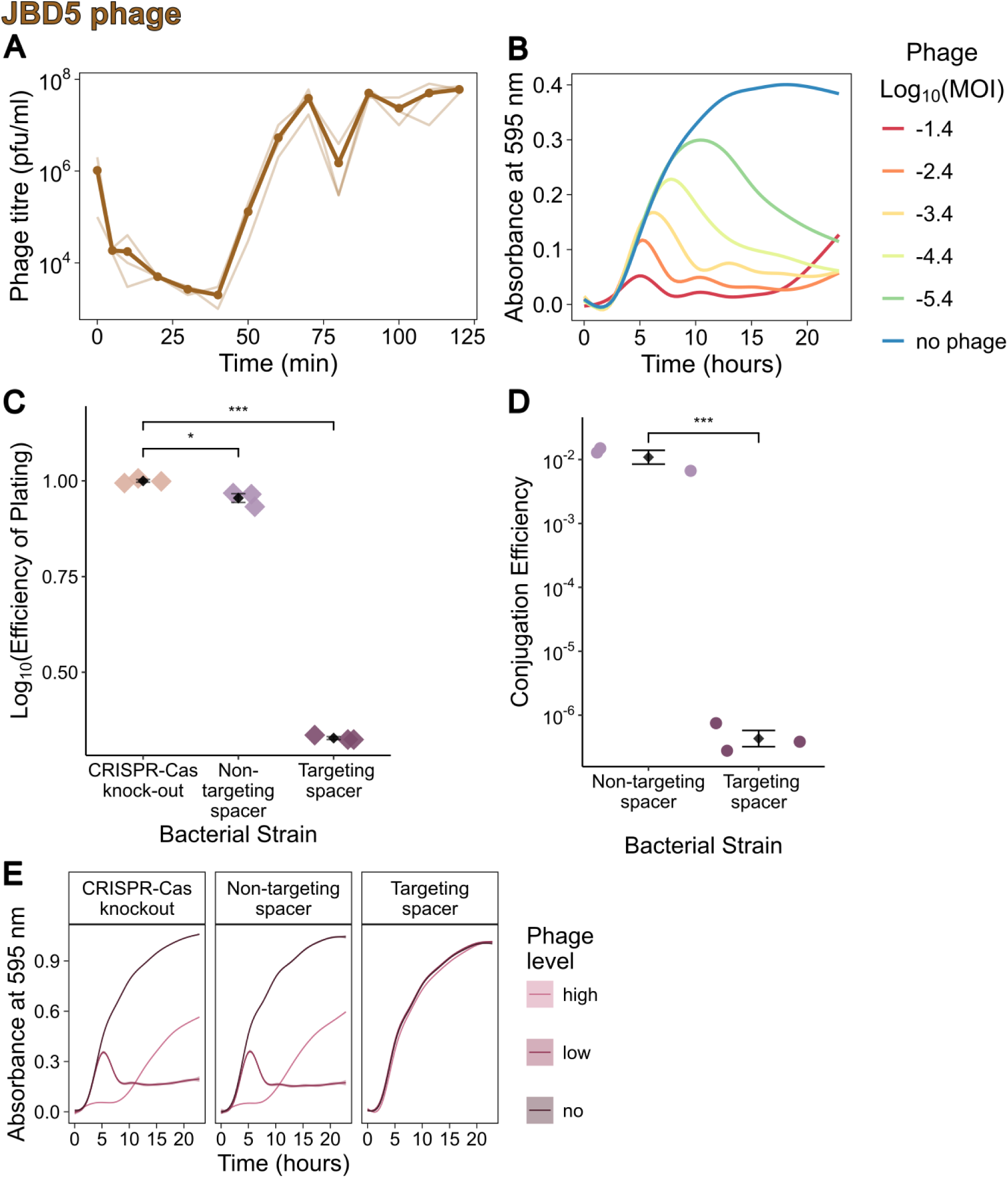
– **JBD5 phage characteristics**. (A) One-step growth curve. Assays were performed at a starting MOI of approximately 0.1. Samples were treated with chloroform to lyse cells, meaning both free phages and mature phages inside cells were measured. The mean (N=3) dots are joined by a bold line with individual samples shown by thin lines. Phages were plated on *P. aeruginosa* wildtype cultures to calculate phage titre (pfu/ml) at each time point. (B) Growth curves in high nutrient liquid media at a range of phage MOIs to calculate virulence with *P. aeruginosa* wildtype. Smoothened mean (N=3) absorbance at OD_595_ lines are coloured rainbow scale red to blue by decreasing log transformed MOI across over 23 hours. (C) Efficiency of plating experiment to test interference by the synthetic CRISPR-Cas system on solid media. Black diamonds denote the mean (N=3), with individual replicate denoted by diamonds coloured by strain type. Error bars give the standard error interval. The CRISPR-Cas knock-out strain is used as the reference strain when calculating efficiency of plating. Statistical testing are pairwise t-tests (* p<0.5, *** p<0.001). On the non-targeting spacer strain the phage grew similarly to the CRISPR-Cas knock-out strain, although there was statistical significance the magnitude of the different in phage titre was small. Whereas the strain with a targeting spacer led to a large and significant decrease in plated phage titre. (D) Conjugation efficiency experiment to validate the phage targeting spacer. The region of the phage genome containing the protospacer targeted by spacer 1 in the synthetic CRISPR array was incorporated into a plasmid. Black diamonds denote the mean (N=3), with individual replicate denoted by dots coloured by strain type. Error bars give the standard error interval. Compared to the non-targeting spacer strain, there was a significant decreased in the proportion of the cell population that took up the protospacer-containing plasmid for the targeting spacer strain (t -test, *** p<0.001). (E) Growth curves for each strain – CRISPR-Cas knock-out, non-targeting spacer and targeting spacer strains in the presence of no phage, a low level of phage (10^4^ pfu/ml), or a higher level of phage (10^6^ pfu/ml). Smoothened mean (N=6) OD_595_ absorbance lines were plotted, coloured by phage level, over a 23-hour growth cycle in nutrient rich liquid media. Each line has ± standard error shaded around the mean line. When *P. aeruginosa* possess the synthetic CRISPR-Cas system with a phage targeting spacer similar growth dynamics are seen with and without phages present.

**Figure S7.**
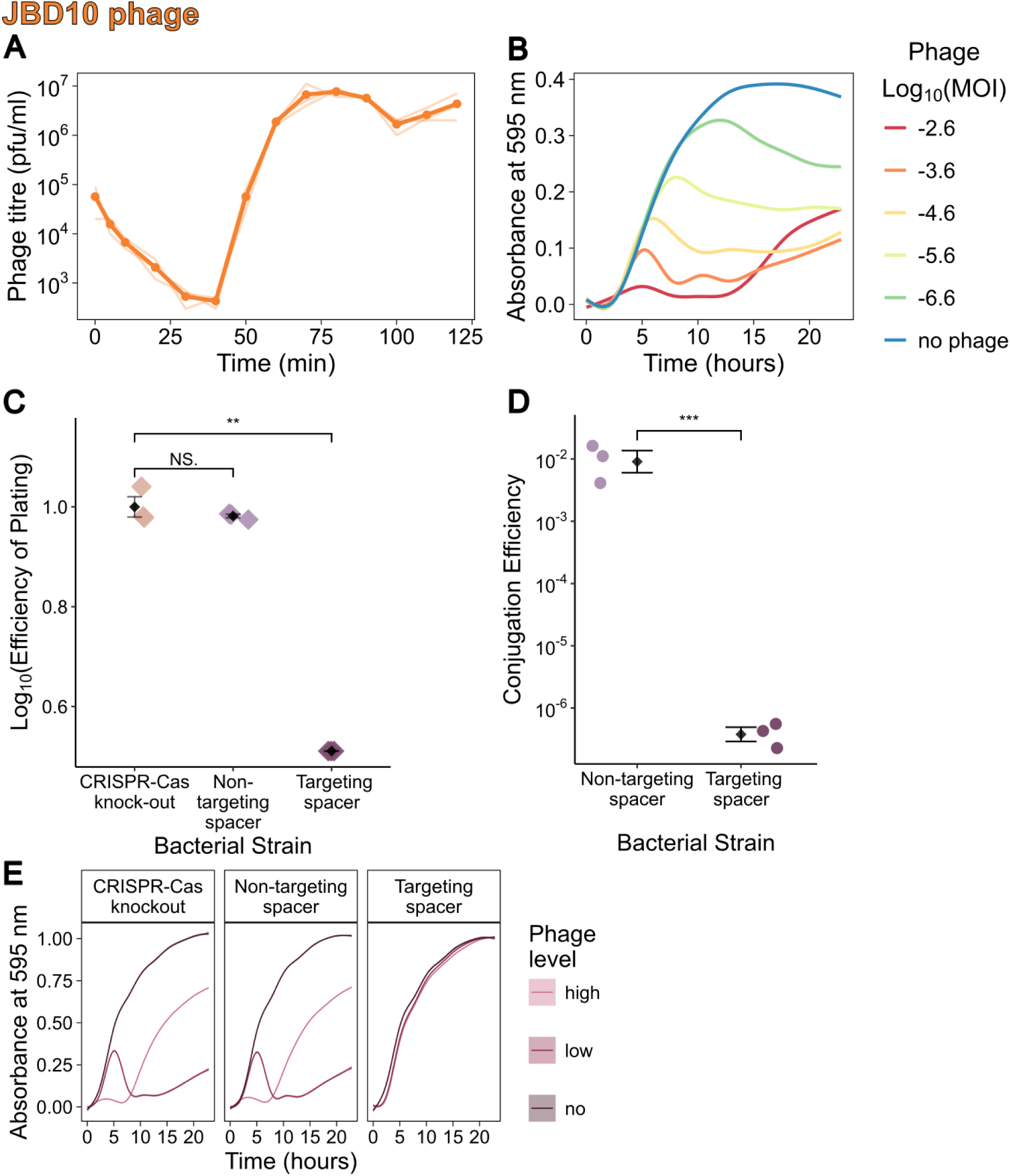
– **JBD10 phage characteristics**. (A) One-step growth curve. Assays were performed at a starting MOI of approximately 0.1. Samples were treated with chloroform to lyse cells, meaning both free phages and mature phages inside cells were measured. The mean (N=3) dots are joined by a bold line with individual samples shown by thin lines. Phage were plated on *P. aeruginosa* wildtype cultures to calculate phage titre (pfu/ml) at each time point. (B) Growth curves in high nutrient liquid media at a range of phage MOIs to calculate virulence with *P. aeruginosa* wildtype. Smoothened mean (N=3) absorbance at OD_595_ lines are coloured rainbow scale red to blue by decreasing log transformed MOI across over 23 hours. (C) Efficiency of plating experiment to test interference by the synthetic CRISPR-Cas system on solid media. Black diamonds denote the mean (N=3), with individual replicate denoted by diamonds coloured by strain type. Error bars give the standard error interval. The CRISPR-Cas knock-out strain is used as the reference strain when calculating efficiency of plating. Statistical testing are pairwise t-tests (NS. non-significant, ** p<0.01). On the non-targeting spacer strain the phage grew similarly to the CRISPR-Cas knock-out strain, whereas the strain with a targeting spacer led to a significant decrease in plated phage titre. (D) Conjugation efficiency experiment to validate the phage targeting spacer. The region of the phage genome containing the protospacer targeted by spacer 1 in the synthetic CRISPR array was incorporated into a plasmid. Black diamonds denote the mean (N=3), with individual replicate denoted by dots coloured by strain type. Error bars give the standard error interval. Compared to the non-targeting spacer strain, there was a significant decreased in the proportion of the cell population that took up the protospacer-containing plasmid for the targeting spacer strain (t -test, *** p<0.001). (E) Growth curves for each strain – CRISPR-Cas knock-out, non-targeting spacer and targeting spacer strains in the presence of no phage, a low level of phage (10^4^ pfu/ml), or a higher level of phage (10^6^ pfu/ml). Smoothened mean (N=6) OD_595_ absorbance lines were plotted, coloured by phage level, over a 23-hour growth cycle in nutrient rich liquid media. Each line has ± standard error shaded around the mean line. When *P. aeruginosa* possess the synthetic CRISPR-Cas system with a phage targeting spacer similar growth dynamics are seen with and without phages present.

**Figure S8.**
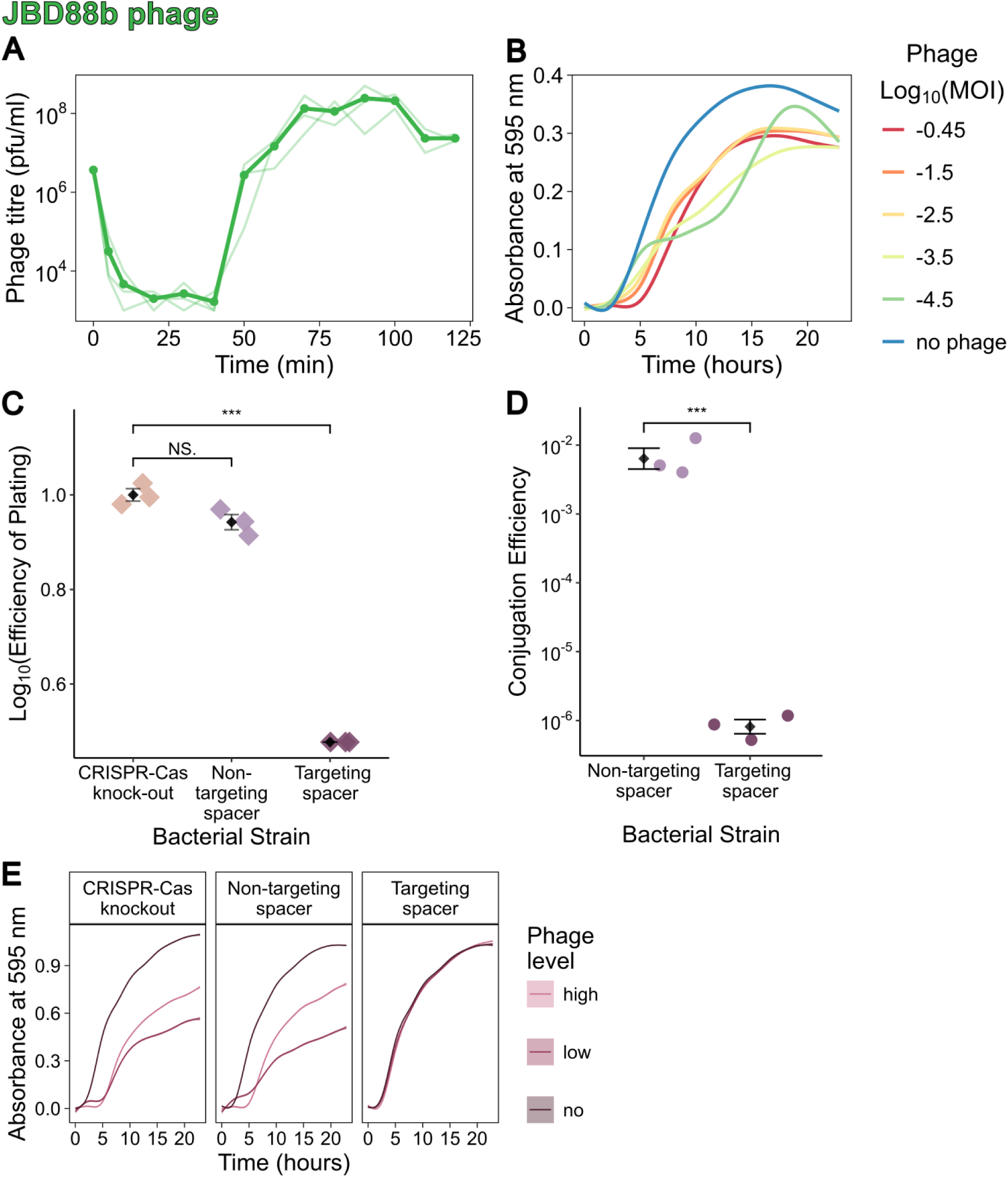
– **JBD88b phage characteristics**. (A) One-step growth curve. Assays were performed at a starting MOI of approximately 0.1. Samples were treated with chloroform to lyse cells, meaning both free phages and mature phages inside cells were measured. The mean (N=3) dots are joined by a bold line with individual samples shown by thin lines. Phages were plated on *P. aeruginosa* wildtype cultures to calculate phage titre (pfu/ml) at each time point. (B) Growth curves in high nutrient liquid media at a range of phage MOIs to calculate virulence with *P. aeruginosa* wildtype. Smoothened mean (N=3) absorbance at OD_595_ lines are coloured rainbow scale red to blue by decreasing log transformed MOI across over 23 hours. (C) Efficiency of plating experiment to test interference by the synthetic CRISPR-Cas system on solid media. Black diamonds denote the mean (N=3), with individual replicate denoted by diamonds coloured by strain type. Error bars give the standard error interval. The CRISPR-Cas knock-out strain is used as the reference strain when calculating efficiency of plating. Statistical testing are pairwise t-tests (NS. non-significant, *** p<0.001). On the non-targeting spacer strain the phage grew similarly to the CRISPR-Cas knock-out strain, whereas the strain with a targeting spacer led to a significant decrease in plated phage titre. (D) Conjugation efficiency experiment to validate the phage targeting spacer. The region of the phage genome containing the protospacer targeted by spacer 1 in the synthetic CRISPR array was incorporated into a plasmid. Black diamonds denote the mean (N=3), with individual replicate denoted by dots coloured by strain type. Error bars give the standard error interval. Compared to the non-targeting spacer strain, there was a significant decreased in the proportion of the cell population that took up the protospacer-containing plasmid for the targeting spacer strain (t -test, *** p<0.001). (E) Growth curves for each strain – CRISPR-Cas knock-out, non-targeting spacer and targeting spacer strains in the presence of no phage, a low level of phage (10^4^ pfu/ml), or a higher level of phage (10^6^ pfu/ml). Smoothened mean (N=6) OD_595_ absorbance lines were plotted, coloured by phage level, over a 23-hour growth cycle in nutrient rich liquid media. Each line has ± standard error shaded around the mean line. When *P. aeruginosa* possess the synthetic CRISPR-Cas system with a phage targeting spacer similar growth dynamics are seen with and without phages present.

**Figure S9.**
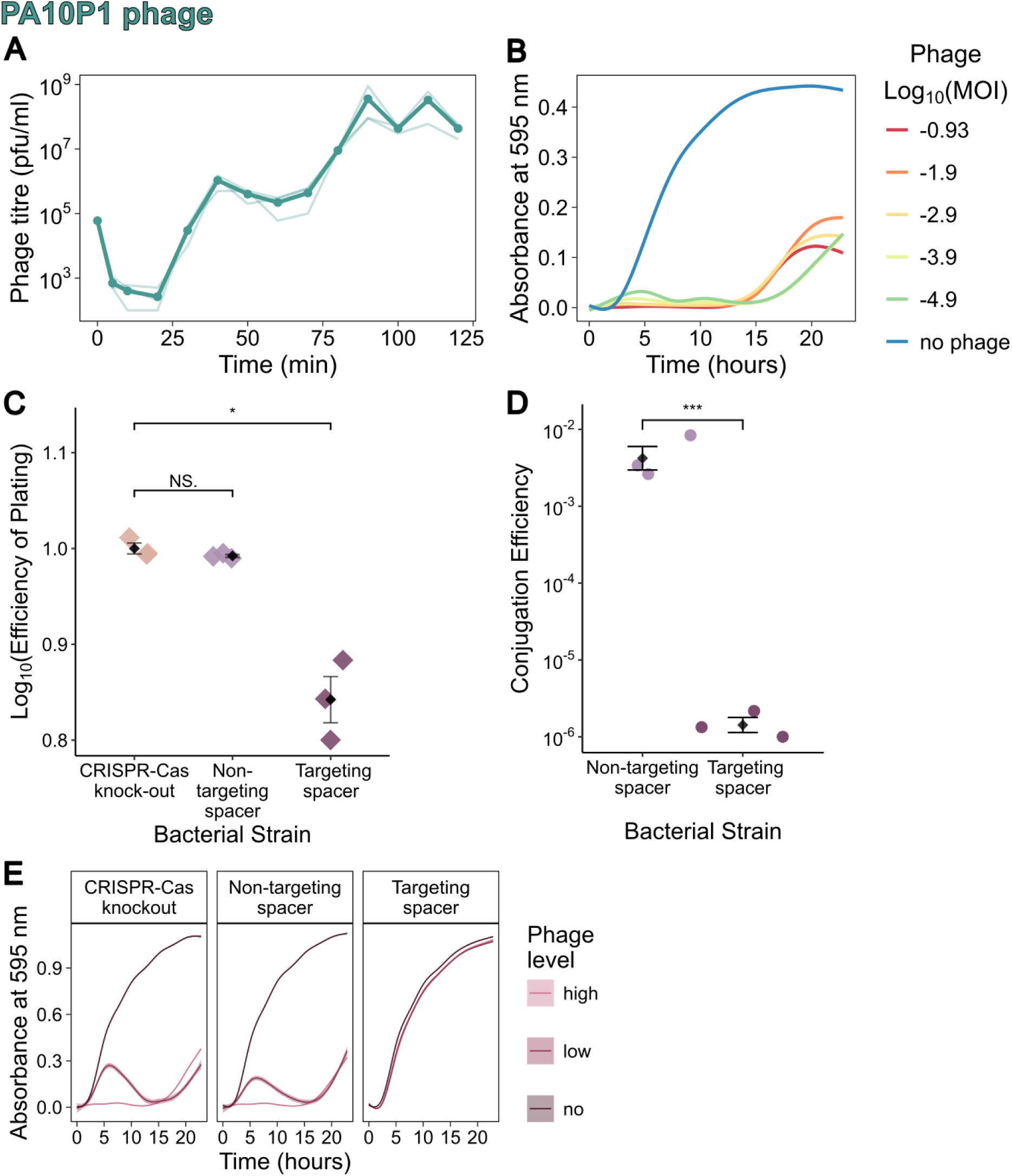
– **PA10P1 phage characteristics**. (A) One-step growth curve. Assays were performed at a starting MOI of approximately 0.1. Samples were treated with chloroform to lyse cells, meaning both free phages and mature phages inside cells were measured. The mean (N=3) dots are joined by a bold line with individual samples shown by thin lines. Phages were plated on *P. aeruginosa* wildtype cultures to calculate phage titre (pfu/ml) at each time point. (B) Growth curves in high nutrient liquid media at a range of phage MOIs to calculate virulence with *P. aeruginosa* wildtype. Smoothened mean (N=3) absorbance at OD_595_ lines are coloured rainbow scale red to blue by decreasing log transformed MOI across over 23 hours. (C) Efficiency of plating experiment to test interference by the synthetic CRISPR-Cas system on solid media. Black diamonds denote the mean (N=3), with individual replicate denoted by diamonds coloured by strain type. Error bars give the standard error interval. The CRISPR-Cas knock-out strain is used as the reference strain when calculating efficiency of plating. Statistical testing are pairwise t-tests (NS. non-significant, * p<0.05). On the non-targeting spacer strain the phage grew similarly to the CRISPR-Cas knock-out strain, whereas the strain with a targeting spacer led to a small but significant decrease in plated phage titre. There was also a large decrease in plaque size on the phage targeting spacer strain (data not shown). (D) Conjugation efficiency experiment to validate the phage targeting spacer. The region of the phage genome containing the protospacer targeted by spacer 1 in the synthetic CRISPR array was incorporated into a plasmid. Black diamonds denote the mean (N=3), with individual replicate denoted by dots coloured by strain type. Error bars give the standard error interval. Compared to the non-targeting spacer strain, there was a significant decreased in the proportion of the cell population that took up the protospacer-containing plasmid for the targeting spacer strain (t -test, *** p<0.001). (E) Growth curves for each strain – CRISPR-Cas knock-out, non-targeting spacer and targeting spacer strains in the presence of no phage, a low level of phage (10^4^ pfu/ml), or a higher level of phage (10^6^ pfu/ml). Smoothened mean (N=6) OD_595_ absorbance lines were plotted, coloured by phage level, over a 23-hour growth cycle in nutrient rich liquid media. Each line has ± standard error shaded around the mean line. When *P. aeruginosa* possess the synthetic CRISPR-Cas system with a phage targeting spacer similar growth dynamics are seen with and without phages present.

**Figure S10.**
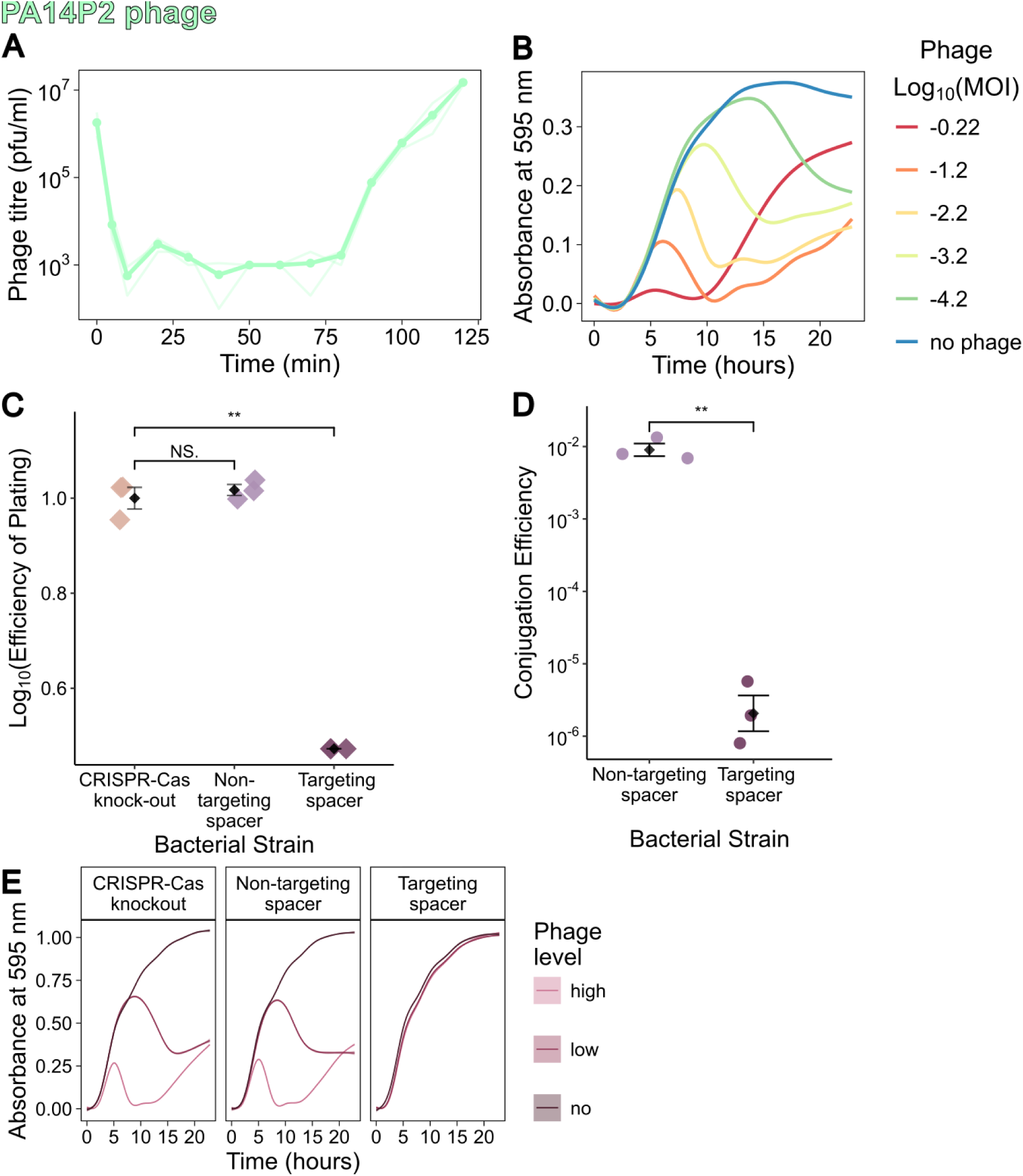
– **PA14P2 phage characteristics**. (A) One-step growth curve. Assays were performed at a starting MOI of approximately 0.1. Samples were treated with chloroform to lyse cells, meaning both free phages and mature phages inside cells were measured. The mean (N=3) dots are joined by a bold line with individual samples shown by thin lines. Phages were plated on *P. aeruginosa* csy3::LacZ cultures to calculate phage titre (pfu/ml) at each time point. (B) Growth curves in high nutrient liquid media at a range of phage MOIs to calculate virulence with *P. aeruginosa* csy3::LacZ. Smoothened mean (N=3) absorbance at OD_595_ lines are coloured rainbow scale red to blue by decreasing log transformed MOI across over 23 hours. (C) Efficiency of plating experiment to test interference by the synthetic CRISPR-Cas system on solid media. Black diamonds denote the mean (N=3), with individual replicate denoted by diamonds coloured by strain type. Error bars give the standard error interval. The CRISPR-Cas knock-out strain is used as the reference strain when calculating efficiency of plating. Statistical testing are pairwise t-tests (NS. non-significant, ** p<0.01). On the non-targeting spacer strain the phage grew similarly to the CRISPR-Cas knock-out strain, whereas the strain with a targeting spacer led to a significant decrease in plated phage titre. (D) Conjugation efficiency experiment to validate the phage targeting spacer. The region of the phage genome containing the protospacer targeted by spacer 1 in the synthetic CRISPR array was incorporated into a plasmid. Black diamonds denote the mean (N=3), with individual replicate denoted by dots coloured by strain type. Error bars give the standard error interval. Compared to the non-targeting spacer strain, there was a significant decreased in the proportion of the cell population that took up the protospacer-containing plasmid for the targeting spacer strain (t -test, ** p<0.01). (E) Growth curves for each strain – CRISPR-Cas knock-out, non-targeting spacer and targeting spacer strains in the presence of no phage, a low level of phage (10^4^ pfu/ml), or a higher level of phage (10^6^ pfu/ml). Smoothened mean (N=6) OD_595_ absorbance lines were plotted, coloured by phage level, over a 23-hour growth cycle in nutrient rich liquid media. Each line has ± standard error shaded around the mean line. When *P. aeruginosa* possess the synthetic CRISPR-Cas system with a phage targeting spacer similar growth dynamics are seen with and without phages present.

**Figure S11.**
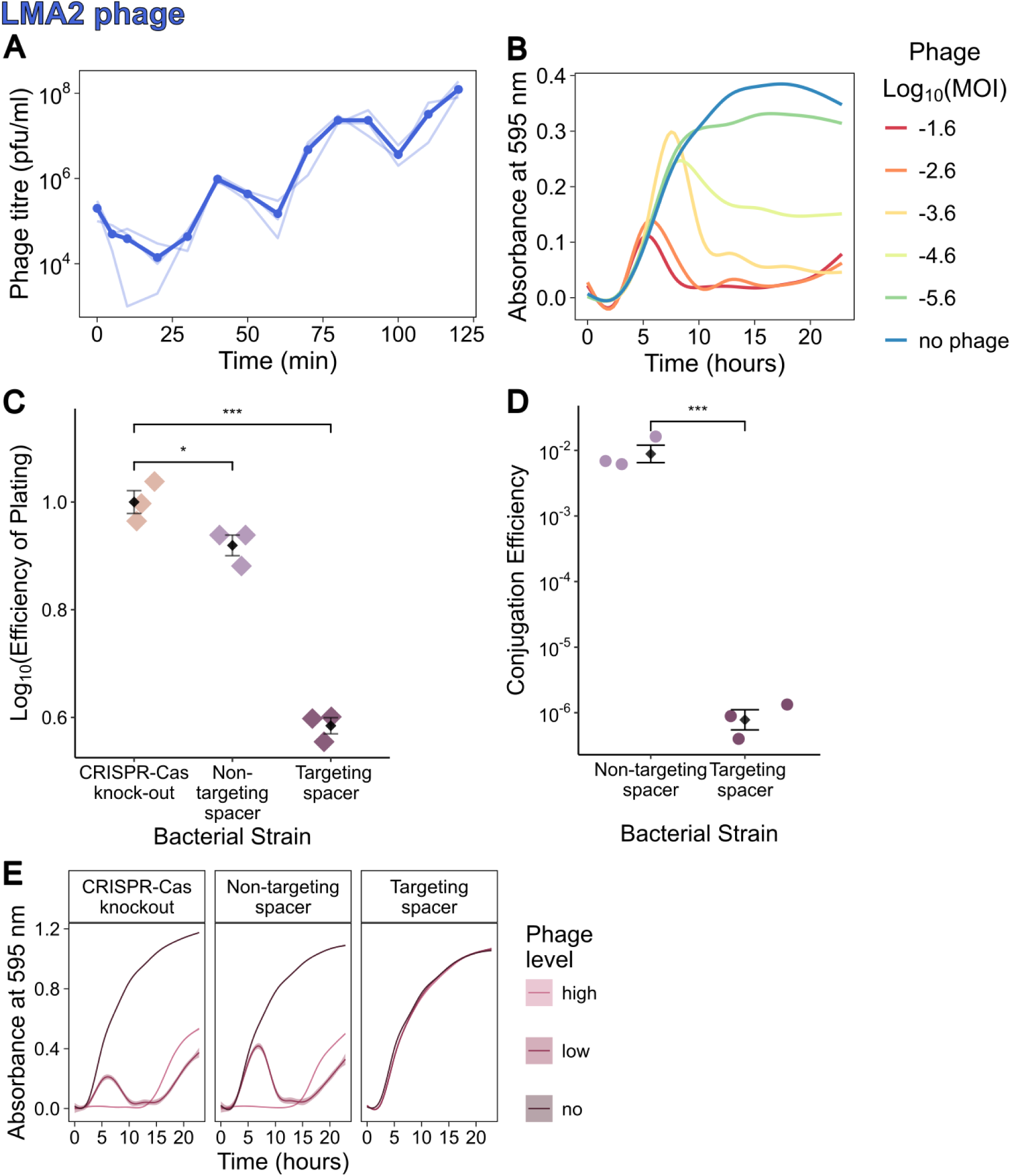
– **LMA2 phage characteristics**. (A) One-step growth curve. Assays were performed at a starting MOI of approximately 0.1. Samples were treated with chloroform to lyse cells, meaning both free phages and mature phages inside cells were measured. The mean (N=3) dots are joined by a bold line with individual samples shown by thin lines. Phages were plated on *P. aeruginosa* wildtype cultures to calculate phage titre (pfu/ml) at each time point. (B) Growth curves in high nutrient liquid media at a range of phage MOIs to calculate virulence with *P. aeruginosa* wildtype. Smoothened mean (N=3) absorbance at OD_595_ lines are coloured rainbow scale red to blue by decreasing log transformed MOI across over 23 hours. (C) Efficiency of plating experiment to test interference by the synthetic CRISPR-Cas system on solid media. Black diamonds denote the mean (N=3), with individual replicate denoted by diamonds coloured by strain type. Error bars give the standard error interval. The CRISPR-Cas knock-out strain is used as the reference strain when calculating efficiency of plating. Statistical testing are pairwise t-tests (* p<0.5, *** p<0.001). On the non-targeting spacer strain the phage grew similarly to the CRISPR-Cas knock-out strain, although there was statistical significance the magnitude of the different in phage titre was small. On the non-targeting spacer strain the phage grew similarly to the CRISPR-Cas knock-out strain, whereas the strain with a targeting spacer led to a significant decrease in plated phage titre. (D) Conjugation efficiency experiment to validate the phage targeting spacer. The region of the phage genome containing the protospacer targeted by spacer 1 in the synthetic CRISPR array was incorporated into a plasmid. Black diamonds denote the mean (N=3), with individual replicate denoted by dots coloured by strain type. Error bars give the standard error interval. Compared to the non-targeting spacer strain, there was a significant decreased in the proportion of the cell population that took up the protospacer-containing plasmid for the targeting spacer strain (t -test, *** p<0.001). (E) Growth curves for each strain – CRISPR-Cas knock-out, non-targeting spacer and targeting spacer strains in the presence of no phage, a low level of phage (10^4^ pfu/ml), or a higher level of phage (10^6^ pfu/ml). Smoothened mean (N=6) OD_595_ absorbance lines were plotted, coloured by phage level, over a 23-hour growth cycle in nutrient rich liquid media. Each line has ± standard error shaded around the mean line. When *P. aeruginosa* possess the synthetic CRISPR-Cas system with a phage targeting spacer similar growth dynamics are seen with and without phages present.

**Figure S12.**
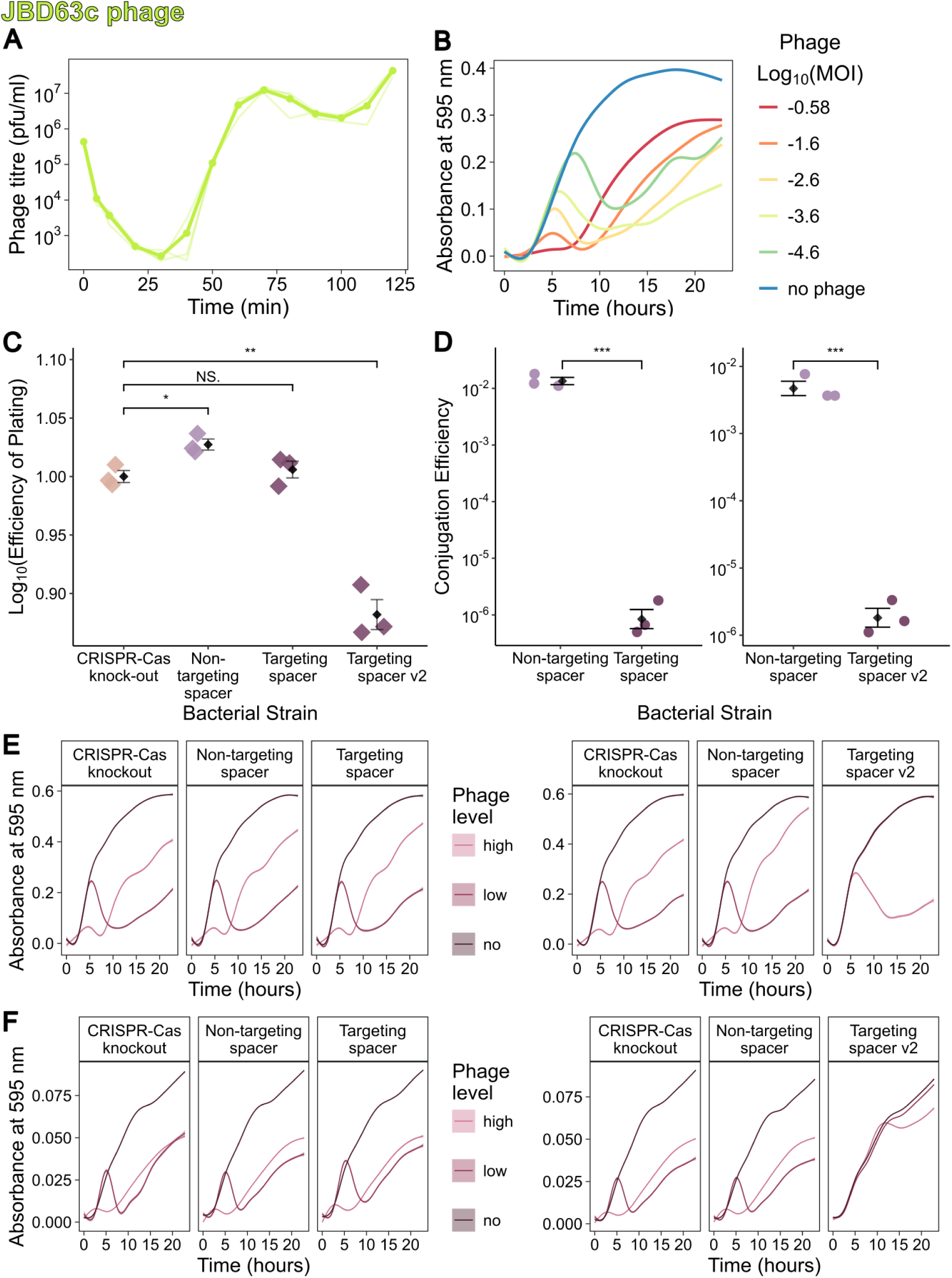
– **JBD63c phage characteristics**. (A) One-step growth curve. Assays were performed at a starting MOI of approximately 0.1. Samples were treated with chloroform to lyse cells, meaning both free phages and mature phages inside cells were measured. The mean (N=3) dots are joined by a bold line with individual samples shown by thin lines. Phages were plated on *P. aeruginosa* wildtype cultures to calculate phage titre (pfu/ml) at each time point. (B) Growth curves in high nutrient liquid media at a range of phage MOIs to calculate virulence with *P. aeruginosa* wildtype. Smoothened mean (N=3) absorbance at OD_595_ lines are coloured rainbow scale red to blue by decreasing log transformed MOI across over 23 hours. (C) Efficiency of plating experiment to test interference by the synthetic CRISPR-Cas system on solid media. Black diamonds denote the mean (N=3), with individual replicate denoted by diamonds coloured by strain type. Error bars give the standard error interval. The CRISPR-Cas knock-out strain is used as the reference strain when calculating efficiency of plating. Statistical testing are pairwise t-tests (NS. non-significant, * p<0.5, ** p<0.01). On the non-targeting spacer strain the phage grew similarly to the CRISPR-Cas knock-out strain, although the difference was significant the magnitude was small. Between the two versions of the targeting spacer strain, the first version with a targeting spacer did now show any difference in efficiency of plating compared to the CRISPR-Cas knock-out strain. Whereas for the targeting spacer version 2 strain there was a modest, but significant, decrease in plated phage titre. (D) Conjugation efficiency experiment to validate the phage targeting spacer. The region of the phage genome containing the protospacer targeted by spacer 1 in the synthetic CRISPR array of each version of the targeting spacer strain version incorporated into each plasmid. Black diamonds denote the mean (N=3), with individual replicate denoted by dots coloured by strain type. Error bars give the standard error interval. Compared to the non-targeting spacer strain, there was a significant decreased in the proportion of the cell population that took up the protospacer-containing plasmid for the targeting spacer strain for both versions of the spacer (t -test, *** p<0.001). (E) High nutrient growth curves for each strain – CRISPR-Cas knock-out, non-targeting spacer and targeting spacer strains in the presence of no phage, a low level of phage (10^4^ pfu/ml), or a higher level of phage (10^6^ pfu/ml). Smoothened mean (N=6) OD_595_ absorbance lines were plotted, coloured by phage level, over a 23-hour growth cycle. Identical experiments were repeated for both versions of the phage targeting spacer strain. Each line has ± standard error shaded around the mean line. When *P. aeruginosa* possess the synthetic CRISPR-Cas system with one version of the phage targeting spacer similar growth dynamics are seen compared to the non-targeting and CRISPR-Cas knockout strains. However, with version 2 of the phage targeting spacer strain, growth is no longer affected by a low level of phage, and growth is improved when exposed to a high dose of phage compared to the non-targeting spacer and CRISPR-Cas knock-out strains. (F) Low nutrient growth curves for each strain exposed the same phage concentrations used in the high nutrient experiment. Smoothened mean (N=3) OD_595_ absorbance lines were plotted, coloured by phage level, over a 23-hour growth cycle. Identical experiments were repeated for both versions of the phage targeting spacer strain. Each line has ± standard error shaded around the mean line. For the first version of the phage targeting spacer, all strains showed the same growth dynamics in the presence of phages. Whereas, for the version 2 of the phage targeting spacer strain, showed the same growth dynamics in the no phage and low-level phage conditions, and similar levels of growth in the high-level phage condition.

**Figure S13.**
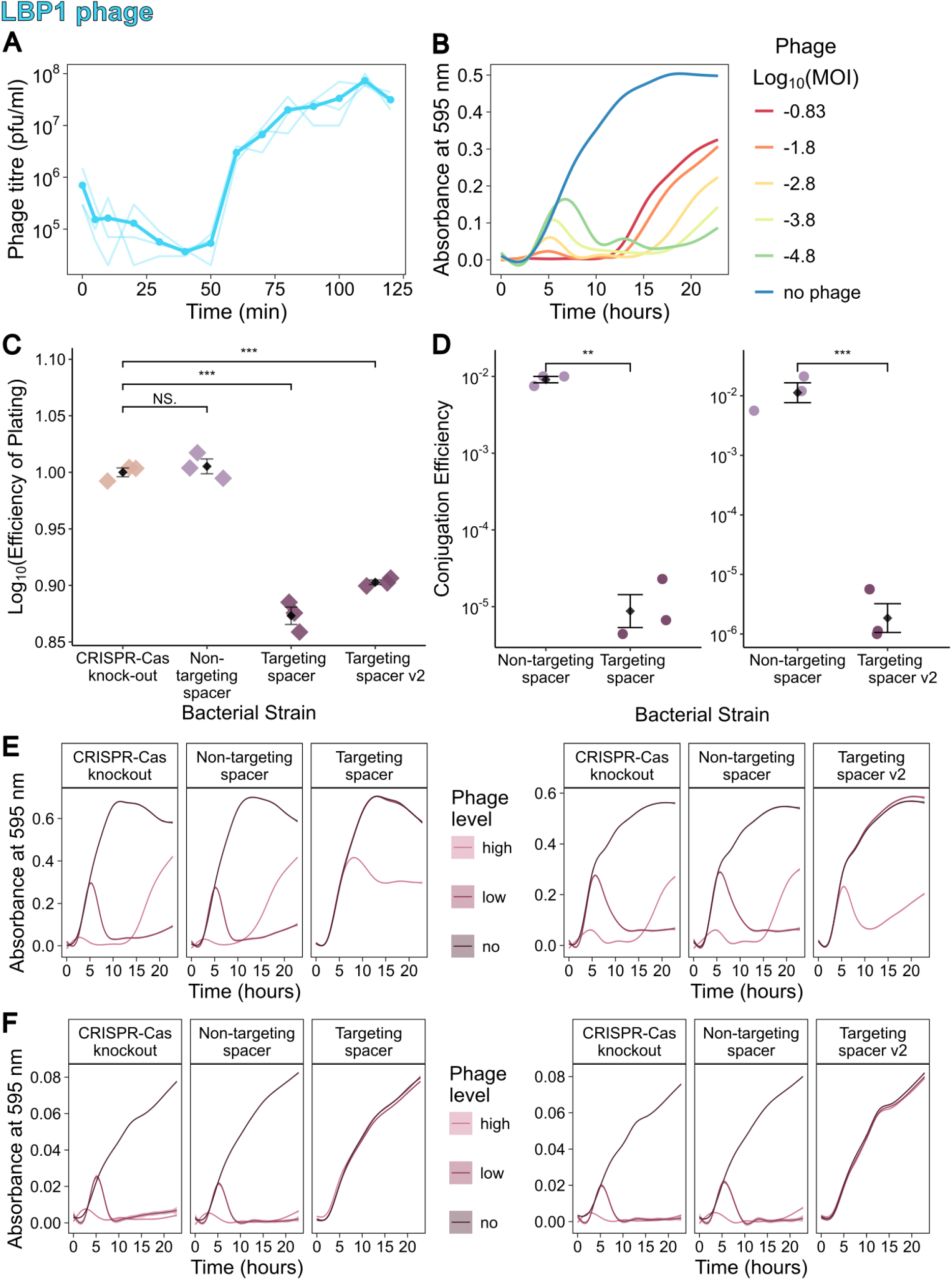
– **LBP1 phage characteristics**. (A) One-step growth curve. Assays were performed at a starting MOI of approximately 0.1. Samples were treated with chloroform to lyse cells, meaning both free phages and mature phages inside cells were measured. The mean (N=3) dots are joined by a bold line with individual samples shown by thin lines. Phages were plated on *P. aeruginosa* wildtype cultures to calculate phage titre (pfu/ml) at each time point. (B) Growth curves in high nutrient liquid media at a range of phage MOIs to calculate virulence with *P. aeruginosa* wildtype. Smoothened mean (N=3) absorbance at OD_595_ lines are coloured rainbow scale red to blue by decreasing log transformed MOI across over 23 hours. (C) Efficiency of plating experiment to test interference by the synthetic CRISPR-Cas system on solid media. Black diamonds denote the mean (N=3), with individual replicate denoted by diamonds coloured by strain type. Error bars give the standard error interval. The CRISPR-Cas knock-out strain is used as the reference strain when calculating efficiency of plating. Statistical testing are pairwise t-tests (NS. non-significant, *** p<0.001). On the non-targeting spacer strain the phage grew similarly to the CRISPR-Cas knock-out strain. For both versions of the phage targeting spacer strains there was a modest, but significant, decrease in plated phage titre. (D) Conjugation efficiency experiment to validate the phage targeting spacer. The region of the phage genome containing the protospacer targeted by spacer 1 in the synthetic CRISPR array of each version of the targeting spacer strain version incorporated into each plasmid. Black diamonds denote the mean (N=3), with individual replicate denoted by dots coloured by strain type. Error bars give the standard error interval. Compared to the non-targeting spacer strain, there was a significant decreased in the proportion of the cell population that took up the protospacer-containing plasmid for the targeting spacer strain for both versions of the spacer (t -test, ** p<0.01, *** p<0.001). (E) High nutrient growth curves for each strain – CRISPR-Cas knock-out, non-targeting spacer and targeting spacer strains in the presence of no phage, a low level of phage (10^4^ pfu/ml), or a higher level of phage (10^6^ pfu/ml). Smoothened mean (N=6) OD_595_ absorbance lines were plotted, coloured by phage level, over a 23-hour growth cycle. Identical experiments were repeated for both versions of the phage targeting spacer strain. Each line has ± standard error shaded around the mean line. For both strains of the *P. aeruginosa* strain with phage targeting spacers, growth is no longer affected by a low level of phage, and growth is improved when exposed to a high dose of phage compared to the non-targeting spacer and CRISPR-Cas knock-out strains. (F) Low nutrient growth curves for each strain exposed the same phage concentrations used in the high nutrient experiment. Smoothened mean (N=3) OD_595_ absorbance lines were plotted, coloured by phage level, over a 23-hour growth cycle. Identical experiments were repeated for both versions of the phage targeting spacer strain. Each line has ± standard error shaded around the mean line. Both versions of *P. aeruginosa* strains with the phage targeting spacer showed the same growth dynamics irrelevant of no, low or high levels of phage exposure.

**Figure S14.**
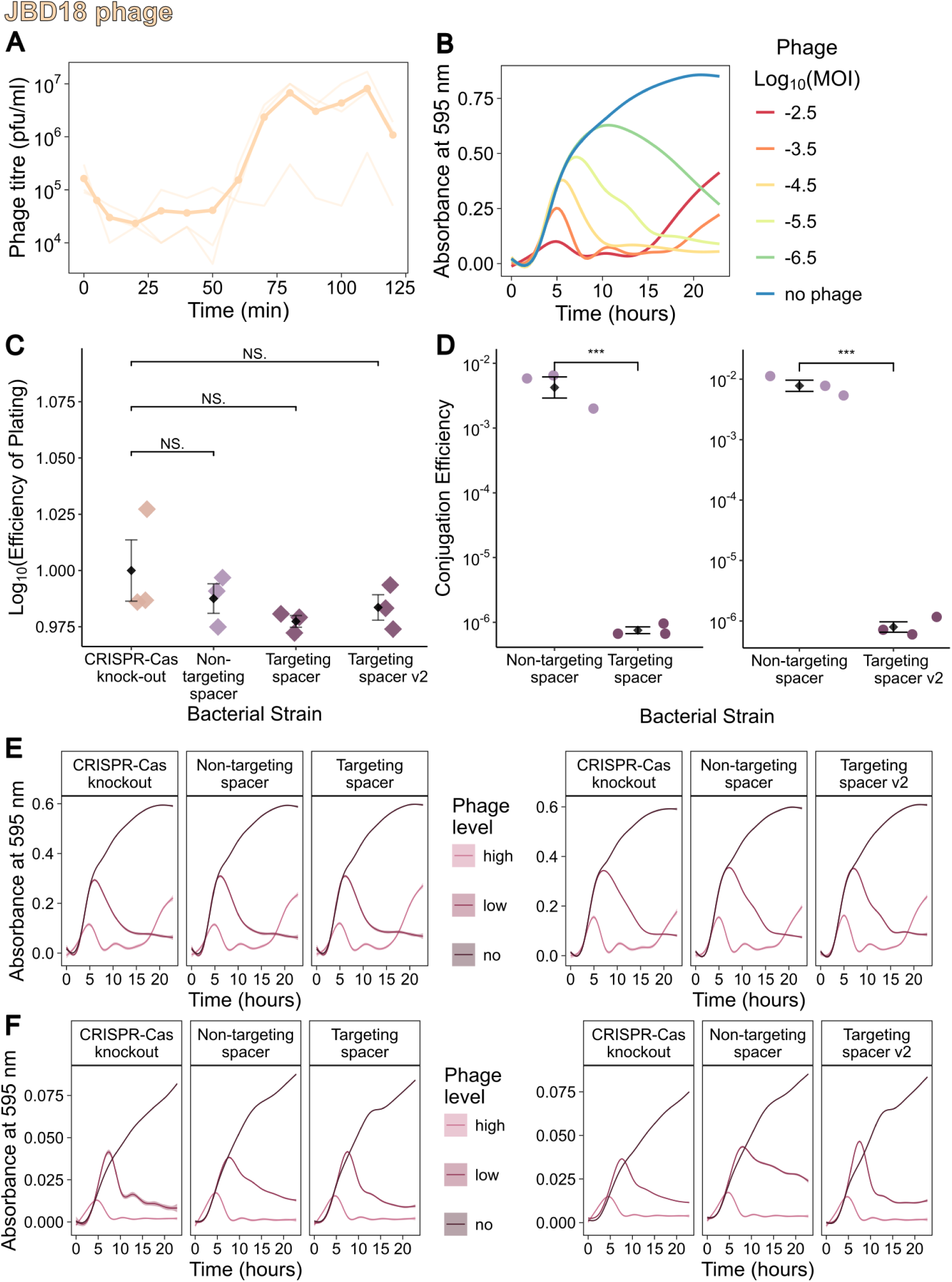
– **JBD18 phage characteristics**. (A) One-step growth curve. Assays were performed at a starting MOI of approximately 0.1. Samples were treated with chloroform to lyse cells, meaning both free phages and mature phages inside cells were measured. The mean (N=3) dots are joined by a bold line with individual samples shown by thin lines. Phages were plated on *P. aeruginosa* wildtype cultures to calculate phage titre (pfu/ml) at each time point. (B) Growth curves in high nutrient liquid media at a range of phage MOIs to calculate virulence with *P. aeruginosa* wildtype. Smoothened mean (N=3) absorbance at OD_595_ lines are coloured rainbow scale red to blue by decreasing log transformed MOI across over 23 hours. (C) Efficiency of plating experiment to test interference by the synthetic CRISPR-Cas system on solid media. Black diamonds denote the mean (N=3), with individual replicate denoted by diamonds coloured by strain type. Error bars give the standard error interval. The CRISPR-Cas knock-out strain is used as the reference strain when calculating efficiency of plating. Statistical testing are pairwise t-tests (NS. non-significant). On the non-targeting spacer strain, and on both phage targeting spacer strains, the phage grew similarly to the CRISPR-Cas knock-out strain. (D) Conjugation efficiency experiment to validate the phage targeting spacer. The region of the phage genome containing the protospacer targeted by spacer 1 in the synthetic CRISPR array of each version of the targeting spacer strain version incorporated into each plasmid. Black diamonds denote the mean (N=3), with individual replicate denoted by dots coloured by strain type. Error bars give the standard error interval. Compared to the non-targeting spacer strain, there was a significant decreased in the proportion of the cell population that took up the protospacer-containing plasmid for the targeting spacer strain for both versions of the spacer (t -test, *** p<0.001). (E) High nutrient growth curves for each strain – CRISPR-Cas knock-out, non-targeting spacer and targeting spacer strains in the presence of no phage, a low level of phage (10^4^ pfu/ml), or a higher level of phage (10^6^ pfu/ml). Smoothened mean (N=6) OD_595_ absorbance lines were plotted, coloured by phage level, over a 23-hour growth cycle. Identical experiments were repeated for both versions of the phage targeting spacer strain. Each line has ± standard error shaded around the mean line. For both versions of the *P. aeruginosa* strain with phage targeting spacers, growth dynamics are the same as those seen in the CRISPR-Cas knock-out and non-targeting spacer strains. (F) Low nutrient growth curves for each strain exposed the same phage concentrations used in the high nutrient experiment. Smoothened mean (N=3) OD_595_ absorbance lines were plotted, coloured by phage level, over a 23-hour growth cycle. Identical experiments were repeated for both versions of the phage targeting spacer strain. Each line has ± standard error shaded around the mean line. For both versions of the *P. aeruginosa* strain with phage targeting spacers, growth dynamics are the same as those seen in the CRISPR-Cas knock-out and non-targeting spacer strains.

**Figure S15.**
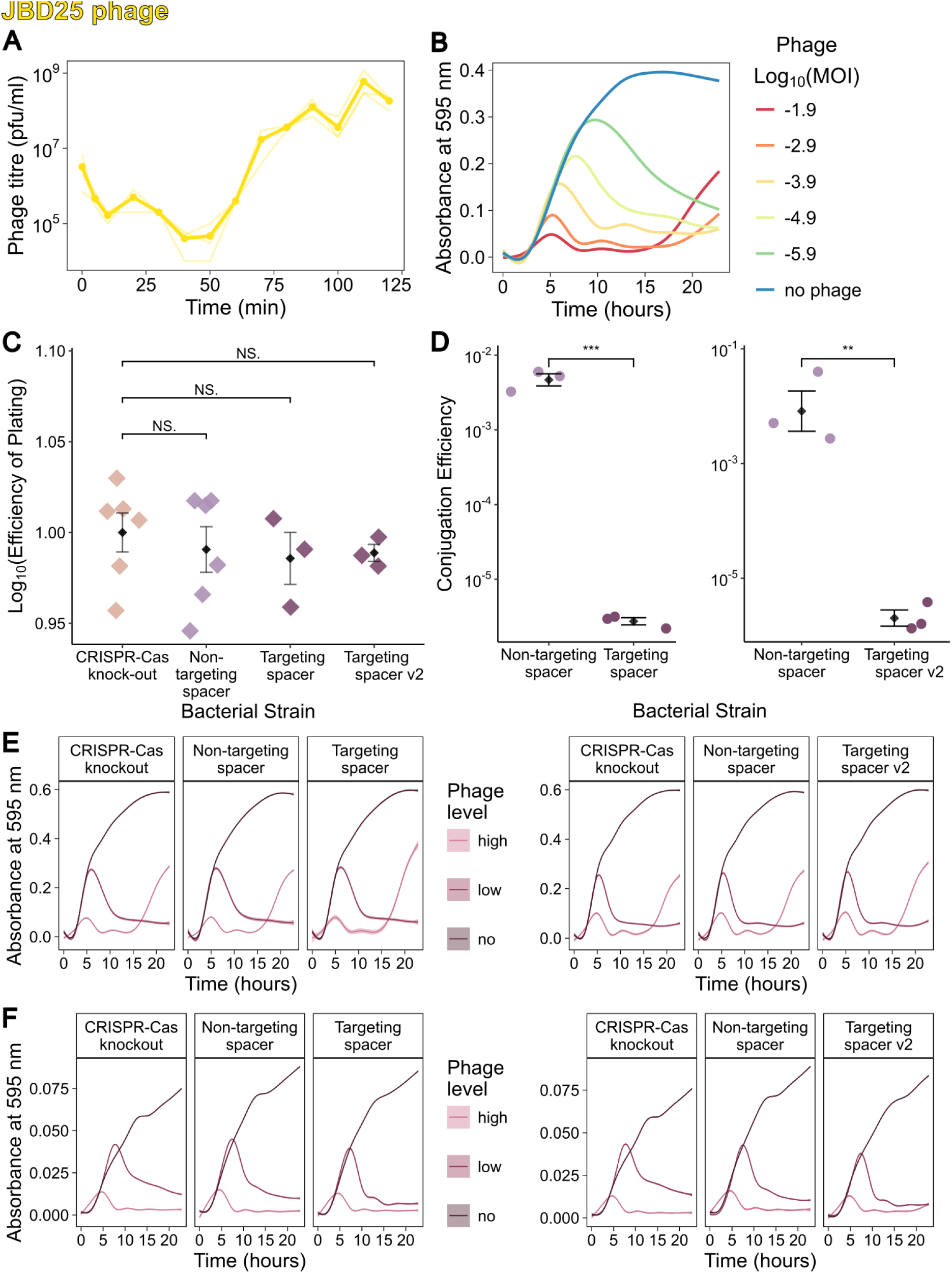
– **JBD25 phage characteristics**. (A) One-step growth curve. Assays were performed at a starting MOI of approximately 0.1. Samples were treated with chloroform to lyse cells, meaning both free phages and mature phages inside cells were measured. The mean (N=3) dots are joined by a bold line with individual samples shown by thin lines. Phages were plated on *P. aeruginosa* wildtype cultures to calculate phage titre (pfu/ml) at each time point. (B) Growth curves in high nutrient liquid media at a range of phage MOIs to calculate virulence with *P. aeruginosa* wildtype. Smoothened mean (N=3) absorbance at OD_595_ lines are coloured rainbow scale red to blue by decreasing log transformed MOI across over 23 hours. (C) Efficiency of plating experiment to test interference by the synthetic CRISPR-Cas system on solid media. Black diamonds denote the mean (N=3), with individual replicate denoted by diamonds coloured by strain type. Error bars give the standard error interval. The CRISPR-Cas knock-out strain is used as the reference strain when calculating efficiency of plating. Statistical testing are pairwise t-tests (NS. non-significant). On the non-targeting spacer strain, and on both phage targeting spacer strains, the phage grew similarly to the CRISPR-Cas knock-out strain. (D) Conjugation efficiency experiment to validate the phage targeting spacer. The region of the phage genome containing the protospacer targeted by spacer 1 in the synthetic CRISPR array of each version of the targeting spacer strain version incorporated into each plasmid. Black diamonds denote the mean (N=3), with individual replicate denoted by dots coloured by strain type. Error bars give the standard error interval. Compared to the non-targeting spacer strain, there was a significant decreased in the proportion of the cell population that took up the protospacer-containing plasmid for the targeting spacer strain for both versions of the spacer (t -test, ** p<0.01, *** p<0.001). (E) High nutrient growth curves for each strain – CRISPR-Cas knock-out, non-targeting spacer and targeting spacer strains in the presence of no phage, a low level of phage (10^4^ pfu/ml), or a higher level of phage (10^6^ pfu/ml). Smoothened mean (N=6) OD_595_ absorbance lines were plotted, coloured by phage level, over a 23-hour growth cycle. Identical experiments were repeated for both versions of the phage targeting spacer strain. Each line has ± standard error shaded around the mean line. For both versions of the *P. aeruginosa* strain with phage targeting spacers, growth dynamics are the same as those seen in the CRISPR-Cas knock-out and non-targeting spacer strains. (F) Low nutrient growth curves for each strain exposed the same phage concentrations used in the high nutrient experiment. Smoothened mean (N=3) OD_595_ absorbance lines were plotted, coloured by phage level, over a 23-hour growth cycle. Identical experiments were repeated for both versions of the phage targeting spacer strain. Each line has ± standard error shaded around the mean line. For both versions of the *P. aeruginosa* strain with phage targeting spacers, growth dynamics are the same as those seen in the CRISPR-Cas knock-out and non-targeting spacer strains.

**Figure S16.**
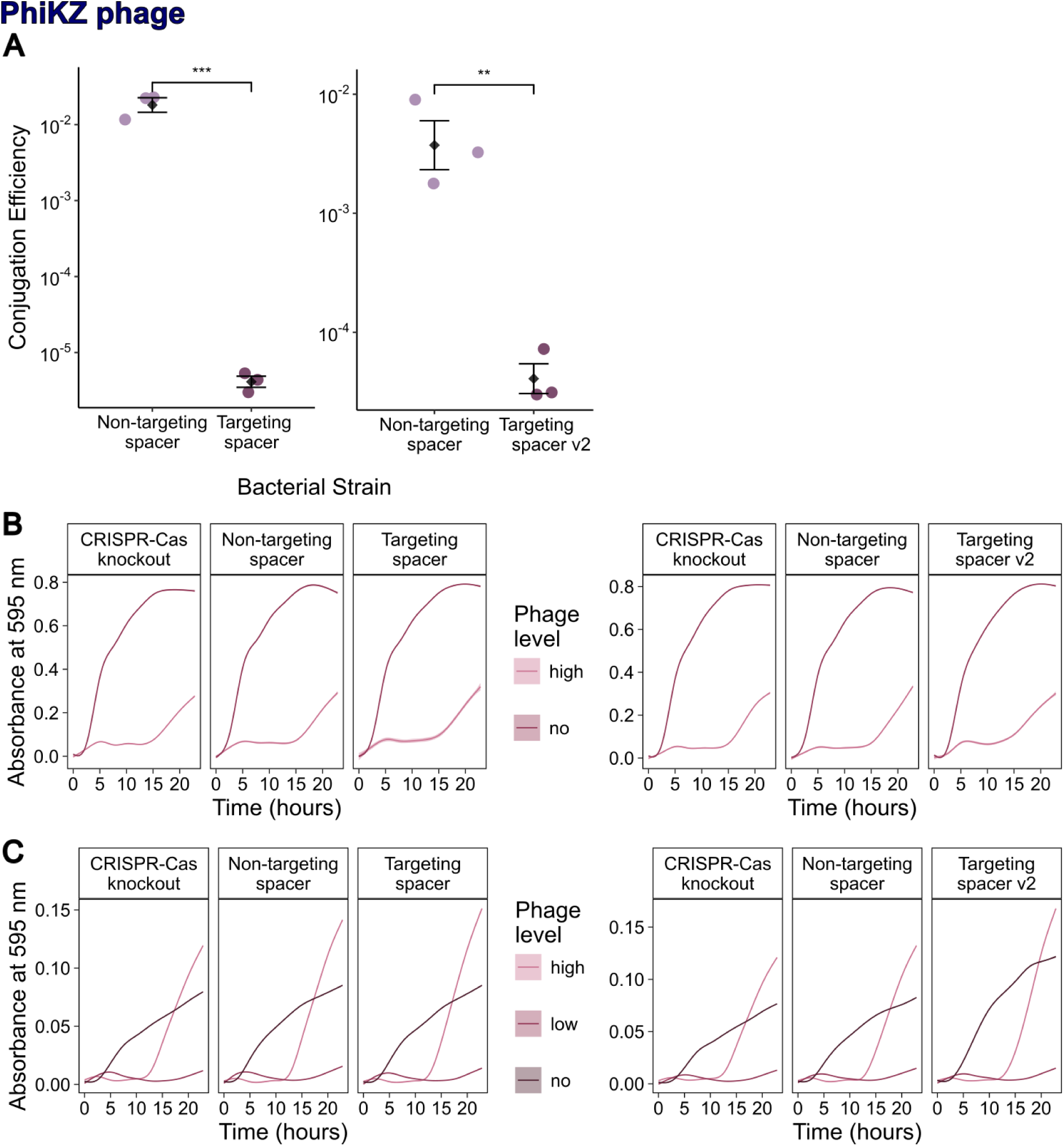
– **PhiKZ phage characteristics**. PhiKZ could not be harvested at a high enough titre to produce a large enough range of MOIs to produce a comparable virulence curve. The plaque morphology of phiKZ on the PA14 strain are small and hazy, making one step growth curves and efficiency of plating experiments challenging to gather accurate titres from. (A) Conjugation efficiency experiment to validate the phage targeting spacer. The region of the phage genome containing the protospacer targeted by spacer 1 in the synthetic CRISPR array of each version of the targeting spacer strain version incorporated into each plasmid.

Black diamonds denote the mean (N=3), with individual replicate denoted by dots coloured by strain type. Error bars give the standard error interval. Compared to the non-targeting spacer strain, there was a significant decreased in the proportion of the cell population that took up the protospacer-containing plasmid for the targeting spacer strain for both versions of the spacer (t -test, ** p<0.01, *** p<0.001). (B) High nutrient growth curves for each strain – CRISPR-Cas knock-out, non-targeting spacer and targeting spacer strains in the presence of no phage, or a higher level of phage (10^6^ pfu/ml) (phages added at a titre of 10^4^ pfu/ml did not produce any effect on cell growth so was excluded from the high nutrient growth curve results). Smoothened mean (N=6) OD_595_ absorbance lines were plotted, coloured by phage level, over a 23-hour growth cycle. Identical experiments were repeated for both versions of the phage targeting spacer strain. Each line has ± standard error shaded around the mean line. For both versions of the *P. aeruginosa* strain with phage targeting spacers, growth dynamics are the same as those seen in the CRISPR-Cas knock-out and non-targeting spacer strains. (C) Low nutrient growth curves for each strain exposed the same phage concentrations used in the high nutrient experiment. Smoothened mean (N=3) OD_595_ absorbance lines were plotted, coloured by phage level, over a 23-hour growth cycle. Identical experiments were repeated for both versions of the phage targeting spacer strain. Each line has ± standard error shaded around the mean line. For both versions of the *P. aeruginosa* strain with phage targeting spacers, growth dynamics are the same as those seen in the CRISPR-Cas knock-out and non-targeting spacer strains.

